# Larger inducible reservoir and higher abundance of exhausted CD8+ T cells in people treated after late versus non-late diagnosis of chronic HIV one year after ART initiation

**DOI:** 10.64898/2026.03.31.715509

**Authors:** Kathryn S. Hensley, Liset de Vries, Tanvir Hossain, Jolieke A.T. van Osch, Raquel Crespo, Michiel Bexkens, Alicja U. Gorska, Cynthia Lungu, Rob A. Gruters, Robert-Jan Palstra, David A.M.C van de Vijver, Jeroen J.A. van Kampen, Peter D. Katsikis, Thibault Mesplède, Shringar Rao, Yvonne M. Mueller, Casper Rokx, Tokameh Mahmoudi

## Abstract

**Background:** Despite effective antiretroviral therapy, HIV-1 remains a global health challenge. Most people living with HIV (PWH) are diagnosed in the chronic stage and around half are diagnosed late globally. Insight into the effect of time of diagnosis and therapy initiation on reservoir dynamics and immune reconstitution within the group with a chronic diagnosis is limited.

**Methods:** In this prospective cohort study, PWH diagnosed during the chronic stage (from Fiebig VI) were stratified into late (<350 CD4+ T cells/mm^3^ or an AIDS-defining illness) or non-late diagnosis subgroups. We analyzed the viral reservoir by IPDA, SQuHIVLa and FISH-Flow, and the immune compartment with the HIV-specific AIM assay and a 45-color spectral flow cytometry panel in the first year after ART initiation.

**Findings:** Although proviral DNA decreased during the first year of ART, the inducible reservoir remained stable. One year after ART initiation, exhausted CD8+ T cell abundance correlated significantly with CD4+ T cell count pre-ART. PWH with a late diagnosis had a significantly higher inducible reservoir and lower CD4+ T-cell counts than the non-late HIV diagnosis subgroup. Moreover, the subgroup with a late diagnosis showed higher abundance of exhausted CD8+ T cells, higher expression of activation/exhaustion markers, and lower naïve CD4+ T-cell abundance than the non-late diagnosis subgroup.

**Interpretation:** Our results show that late diagnosis is associated with a persistently higher inducible viral reservoir and impaired immune recovery. These findings underline the importance of early diagnosis and treatment, and rationalize the use of late diagnosis as a covariate in future cure and reservoir studies.

**Clinicaltrials.gov identifier:** NCT04888754

## Introduction

Despite the effectiveness of antiretroviral therapy (ART) in suppressing HIV replication, HIV-1 remains a major global health challenge, with 1.3 million new cases annually and nearly 40 million people affected worldwide (1). ART requires lifelong adherence to prevent viral rebound, posing challenges such as drug toxicity, resistance and high costs (2, 3). Even with effective ART, people living with HIV (PWH) experience higher rates of morbidity and mortality due to persistent inflammation (4). Moreover, stigma and inequitable access to treatment continue to hinder global HIV control, highlighting the need for a cure (5).

The main obstacle to HIV cure is the persistent reservoir of infected long-lived cells that can evade elimination by the immune system (6, 7). A range of molecular mechanisms contribute to the persistence of the HIV reservoir, including transcriptional and post-transcriptional blocks (8–10). The reservoir can express low levels of viral transcripts and occasionally proteins despite suppressive ART (11–14). Comprehensive characterization and quantification of the HIV reservoir and immune function in PWH on suppressive ART is essential for advancing our understanding of viral persistence, particularly in the context of developing cure strategies.

Early initiation of ART limits reservoir seeding, reduces immune activation, and preserves immune function (15–17). Consequently, individuals initiating ART during the acute stage of the infection have become a central focus in HIV cure research (18). However, only 8.4% of HIV-1 diagnoses in Europe are made at the acute stage, and globally this percentage varies between 1 and 20%, depending on the setting (19–21). Most PWH initiate ART during the chronic phase (from Fiebig stage VI), which makes this a particularly relevant population to study (22).

Treatment initiated at the chronic stage of HIV-1 infection has been reported to result in higher total HIV DNA, intact HIV DNA, and cell associated HIV RNA than when treatment is initiated at the acute stage (16, 23–25). Moreover, PWH with a diagnosis established during chronic disease have more T cell activation on ART, both in CD4+ as well as CD8+ T cells (23). When examining differences between chronic and acute diagnosis, the chronic diagnosis group includes PWH diagnosed immediately after Fiebig stage VI as well as those treated at more advanced stages, including people who have developed AIDS. Despite asserted clinical differences between people living these two extremes, there is limited research examining heterogeneity within the chronic diagnosed population.

Clinical parameters allow for the classification of the chronic diagnosed population into late- and non-late diagnosis, with a late diagnosis typically defined by CD4+ T cell counts below 350 cells/mm³ or an AIDS-defining illness (19, 26). Approximately 40 to 50% of PWH worldwide are diagnosed at a the late stage (19, 27, 28), often accompanied by a higher prevalence of opportunistic infections, slower time to viral suppression, and increased mortality (15, 29–33). Therefore, HIV-1 treatment guideline recommendations were adjusted to offer treatment directly after diagnosis instead of waiting for a CD4 T cell count drop or the development of opportunistic infections (31, 34).

Although the profound difference between a non-late and late chronic diagnosis has been well-studied within clinical settings, little is known about potential differences in the viral reservoir between these two subgroups. Similarly, in-depth analysis of immune reconstitution comparing these subgroups is lacking. Often no distinction is made between late- and non-late diagnoses in inclusion of participants in HIV-1 cure trials or in viral reservoir studies. It is unclear how this heterogeneity in the chronically diagnosed group may impact viral reservoir and cure studies.

To address this gap, we evaluate reservoir dynamics and recovery within the immune compartment in a cohort of participants diagnosed and treated at the chronic stage of infection. We established a longitudinal cohort of 35 PWH in the Netherlands diagnosed at the chronic stage, with sampling time points at start of ART (week 0) as well as 24 and 52 weeks after ART initiation (NCT04888754). We used distinct reservoir quantitation technologies to characterize the reservoir at both the (intact) DNA, and inducible RNA and protein levels while the immune compartment was characterized using a 45-color immunophenotyping panel (OMIP-109) (35) as well as the HIV-specific immunity with activation-induced marker (AIM) assay. Using these approaches, we demonstrate that a chronic but non-late diagnosis is associated with a smaller inducible multiple spliced RNA reservoir and lower abundance of exhausted CD8+ effector memory T cells when compared to a late diagnosis. These findings support early diagnosis and rapid treatment initiation, even in the chronic stage of infection. The findings also highlight potential implications for HIV-1 cure trial design, such as participant classification and selection criteria in cure studies.

## Methods

### Study design and participants

We conducted a prospective observational cohort study at the Erasmus University Medical Centre, Rotterdam, the Netherlands. ART naïve adults diagnosed from Fiebig stage VI and who initiated integrase inhibitor-based triple drug ART in routine care were eligible for participation. The Fiebig stage VI was defined as having a fully converted HIV western blot at the time of ART initiation. In total, 35 participants were included.

### Clinical procedures

Participants were included within seven days after starting ART. Visits with phlebotomy sampling occurred at week 0 (inclusion), week 24, and week 52. Optional sampling via leukapheresis occurred at weeks 24 and 52. Clinical data collected at baseline included age, sex assigned at birth, country of birth, and at each visit we evaluated medical history, plasma HIV-RNA, CD4+ T cell counts, and ART/co-medication use.

Based on WHO/ECDC classifications, we defined a person as having received a late diagnosis when they had a CD4+ T cell count below 350 cells/mm^3^ at the time of HIV diagnosis, or if they presented with an AIDS-defining illness regardless of CD4+ T cell count (19, 26). Participant HIV Indicator conditions and comedications used by participants are shown in Table S1. In this study, viral suppression was defined as achieving a plasma HIV-RNA level below 30 copies/ml, which is the lower limit of quantification of the HIV-RNA assay used. All but one participant were suppressed before week 52. One participant showed low level viremia with viral loads of 274 and 168 copies/mL at weeks 24 and 52 respectively. This participant was also treated with immuno-suppressant medication at all timepoints and was excluded from reservoir or immune characterization but included in clinical endpoints (Table S2). One participant had a detectable viral load (34 copies/mL) at week 24 and was suppressed at week 52. Two participants experienced a viral blip: one at week 24 (52 copies/mL) and another at week 52 (79 copies/mL). Not all techniques were possible to be performed on all participants due to subtype mismatches and sample availability (Table S2).

### Diagnostic Laboratory procedures

The detection of HIV antibodies and/or p24 antigen was accomplished using a fourth-generation HIV test (Liaison XL HIV Ab/Ag chemiluminescence immunoassay; Diasorin). In the event of positive results from this fourth-generation HIV test, confirmation was obtained through an HIV immunoblot (INNO-LIA HIV I/II Score from Fujirebio), and an HIV-1 nucleic acid amplification test (NAAT) (Aptima HIV-1 Dx Quant Assay from Hologic). Follow-up HIV-1 RNA monitoring was performed using the same test. Resistance mutation testing was performed for all participants on integrase, protease, and reverse transcriptase. HIV-1 subtype was based on these results.

### PBMC and CD4+ T cell isolation

Peripheral blood mononuclear cells (PBMCs) were isolated using density gradient centrifugation with Ficoll-Hypaque (GE Healthcare Life Sciences) and collected in RPMI- 1640 medium (Life Technologies) supplemented with 3% fetal bovine serum (FBS). The PBMCs were washed three times, frozen in a freezing medium consisting of 90% FBS and 10% dimethyl sulfoxide (DMSO), and stored in liquid nitrogen until further use. Primary CD4+ T cells were isolated from freeze-thawed PBMCs through magnetic separation with EasySep Human CD4+ T cell Enrichment kit (StemCell Technologies, Grenoble, France) according to the manufacturer’s instructions.

### DNA isolation

DNA was extracted from isolated CD4+ T cells using the Puregene Cell Core Kit, with additional precautions taken to prevent DNA shearing. Briefly, 3.5*10^6^ CD4+ T-cells were lysed, then the protein precipitation solution was added, and samples were incubated on ice for fifteen minutes. Samples were centrifuged and the supernatant was collected. DNA was purified by isopropanol precipitation. Samples were never vortexed during resuspension. Purified DNA was digested with Not-I-HF (NEB) and quantified using Qubit.

### Intact proviral DNA assay (IPDA)

The original IPDA was adapted and validated for specificity and sensitivity using qPCR and dPCR (36, 37)(Table S3, supplementary data, Fig. S1A). Reservoir quantification was performed on isolated CD4+ T cells from participants with HIV-1 subtype B virus collected at 24 and 52 weeks after antiretroviral therapy initiation, using the QuantStudio Absolute Q Digital PCR System (ThermoFisher Scientific). Samples were analyzed in triplicate, using 700ng of DNA. Parallel analysis of RPP30 was performed using 7ng of DNA, as previously described(36), to calculate the DNA shearing index and measure input cell number. Data analysis was performed using the QuantStudio Absolute Q Software Version 6.3. Statistical analyses were performed using GraphPad Prism 10.6.1 (GraphPad Software LLC). Paired comparisons were performed using the Wilcoxon signed-rank test.

### FISH-Flow

Fluorescent in-situ hybridisation flow cytometry (FISH-Flow) was performed using the PrimeFlow RNA assay (Thermo Fisher Scientific) following the manufacturer’s instructions and as described before (38, 39). After thawing and resting, CD4+ T cells were stimulated for 18 hours with phorbol 12-myristate 13-acetate (PMA) and ionomycin (Sigma-Aldrich) at a concentration of 100 ng/ml and 1 μg/ml, respectively. After this, CD4+ T cells were stained with fixable viability dye 780 (Thermo Fisher Scientific, 65086514) using a dilution of 1:1000 for 20 minutes at 4 °C. After fixation and permeabilisation, the intracellular Gag protein was stained using the anti-HIV-1 Gag antibodies KC57 RD-1 (Beckman Coulter, 6604665) and 28B7 (Medimabs) at a final dilution of 1:400 and 1:750, respectively, at 4 °C for 1 hour. mRNA was labelled with a set of 40 probe pairs targeting the GagPol region of the viral RNA (catalog number GagPol HIV-1 VF10-10884, Thermo Fisher Scientific) diluted 1:5 in the diluent provided in the kit and hybridised to the target mRNA for 2 hours at 40°C. Samples were washed to remove excess probes and stored overnight in the presence of RNAsin. Signal amplification was realized by incubation of the cells at 40 °C for 1.5 hour with the pre-amplification mix and subsequently with the amplification mix. Next, the amplified signal was labelled with the label probe mix for 1 hour at 40 °C. Samples were acquired on a BD LSRFortessaTM Analyzer. The analysis was performed using the FlowJo V10.6.1 software (Treestar). The gating strategy is detailed in Fig S1B.

### SQuHIVLa assay

The Specific Quantification of Inducible HIV-1 by RT-LAMP (SQuHIVLa) was performed as described previously (40). Briefly, isolated CD4+ T cells were resuspended in RPMI-1640 culture medium supplemented with 10% FBS and 100 µg/mL penicillin-streptomycin. The cells were rested for 5 hours at 37°C in a humidified incubator with 5% CO₂. Subsequently, the cells were stimulated with 100 ng/mL PMA and 1 µg/mL ionomycin for 12 hours at 37°C with 5% CO₂. Following activation, the cells were washed, counted, serially diluted in Phosphate-buffered saline (PBS), and distributed into a 96-well PCR plate containing 15 µL of RT-LAMP master mix. The RT-LAMP reaction was conducted with an initial incubation at 45°C for 60 minutes, followed by continuous amplification at 65°C, with fluorescence measurements taken every 30 seconds over 180 cycles (equivalent to 90 minutes of amplification time, unless stated otherwise). After completing the RT-LAMP reaction, the frequency of cells expressing tat/rev multiply spliced (ms)RNA was calculated using IUPMStats v1.0 online software, using the maximum likelihood method(41).

### AIM assay

T cell responses to HIV were characterized using the activation-induced marker (AIM) assay on cryopreserved PBMCs. After thawing and resting, 1x10^6^ PBMCs were incubated with HIV- 1 B-specific peptide pools Env, Gag, Nef, Pol, and Tat (all 1µg/mL) for 20 hours at 37 °C with 5% CO2. PBMCs were stimulated with an equimolar amount dimethyl sulfoxide (DMSO) as negative control and CMV pp65 as positive control, DMSO background percentage was subtracted from each peptide-stimulated well. PBMCs were stained for 30 minutes at 4°C with the following antibodies and dilutions: anti-CD3PE-eFluor 610 (Clone UCHT1, eBioscience, 1:100), anti-CD4eFluor450 (Clone HI30, eBioscience, 1:50), anti-CD8Alexa Fluor 647 (Clone RPA-T8, BD, 1:50), anti-CD45RABV605(Clone HI100, Biolegend, 1:50), anti- CCR7BB700 (Clone 3D12, BD, 1:50), anti-CD69FITC (Clone FN50, eBioscience, 1:50), anti- CD137/41-BBPE (Clone 4B4-1, Biolegend, 1:50), and anti-OX40BV510 (Clone Ber-ACT35, Biolegend, 1:25). The cells were subsequently stained for 15 minutes at 4°C with LIVE/DEAD staining (Fixable viability dye 780, Thermofisher, 1:1000). After staining the cells were analyzed by flow cytometry (FACSFortessa, BD Biosciences). HIV-reactive T cells were identified as 4-1BB+OX40+ and 4-1BB+CD69+ for CD4+ T cells, and 4-1BB+CD69+ for CD8+ T cells. On average, 250 000 events were measured per condition. The gating strategy is detailed in Fig S1C.

### In-depth immunophenotyping

For in-depth immunophenotyping PBMCs were thawed, washed twice with RPMI + 3% heat-inactivated FBS and stained using a 45-colour flow cytometry panel. Data acquisition was performed on a 5-laser Aurora spectral flow cytometer (Cytek Biosciences, CA). Staining followed the OMIP-109 protocol with minor modifications (25). The live/dead stain was substituted with an Annexin V conjugate (Thermo Fisher Scientific, Cat# A23202) to exclude both dead and apoptotic cells. Additionally, all the buffers were supplemented with 2.5 mM CaCl₂. To improve resolution of the CD20+CD19+ population, anti-CD20 antibody was added as a final step of the sequential stainings for 5 minutes at room temperature, prior to the addition of the full antibody cocktail. Spectral unmixing and compensation checks were conducted using SpectroFlo software (v2.2.0). All analyses were performed within the OMIQ software environment (www.omiq.ai).

### Outcomes

The main study endpoint was the evolution in reservoir size as assessed as the number of cells expressing viral RNA and viral protein in PWH on ART at week 24 and 52 after treatment initiation. Secondary endpoints included reservoir size, inducibility of the reservoir, the evolution of T cell responses and its relation to the reservoir, and evaluation of clinical parameters with reservoir size, activity, and host immune responses.

### Sample size and statistical analysis plan

The sample size calculation, based on preliminary results from SQuHIVLa experiments and studies on CD4+ T cells from PWH (40, 42), estimated the mean frequency of cells capable of expressing multiple spliced HIV-1 RNA to be 18 cells per million CD4+ T cells after PMA/ionomycin stimulation at week 52. To detect a 50% reduction in reservoir size, from 36 cells per million CD4+ T cells at week 24 to 18 cells per million CD4+ T cells (Multiple spliced HIV-1 RNA+) at week 52, a minimum of 13 participants was required. This sample size provides 80% power to detect the specified reduction, assuming a standard deviation of 20 CD4+ T cells (multiple spliced HIV-1 RNA+) per million and an alpha of 0.05. Due to sample availability, not all assays were performed on all donors at all timepoints. Characteristics of each group of donors within each technique were similar to the overall cohort (Table S3).

Categorical data are presented as n (%), and continuous variables are presented as mean (95% CI) or median (IQR), depending on the normality of the data (assessed by visual inspection and/or the Kolmogorov-Smirnov test). Categorical data were compared using Fisher’s exact test. Continuous data were compared using the Mann-Whitney U test (two groups) or Kruskal-Wallis test (three or more groups); paired data used the Wilcoxon signed-rank test, or Friedman test for multiple timepoints. Correlations with continuous pre-ART CD4 count were assessed by Spearman’s rank correlation. The immunophenotyping screen (522 features) was corrected for multiple testing using the Benjamini-Hochberg procedure (FDR < 0.05). PCA was performed on z-scored feature abundances, both across the full panel and aggregated by lineage compartment. Analyses were performed in Python.

## Ethical considerations

The study was performed in accordance with the principles of the Declaration of Helsinki, Good Clinical Practice guidelines, and the Dutch Medical Research Involving Human Subjects Act (WMO). Written informed consent was obtained from all participants. The study was reviewed and approved by the Erasmus MC Medical Ethics Committee (MEC 2020-0165) and was registered at clinicaltrials.gov (NCT04888754).

## Results

### Cohort characteristics, inclusions, and clinical outcomes

To investigate the heterogeneity in reservoir dynamics and immune reconstitution within PWH diagnosed and treated at the chronic stage of HIV-1 infection, 108 people newly diagnosed with chronic HIV were screened, leading to the inclusion of 35 participants in the cohort (Fig. S1D). Most participants included were assigned male at birth (86%) and had an HIV-1 subtype B (69%) (Table S5). Median CD4+ T cell count before ART initiation was 360 cells/mm^3^ (IQR 210-510). Approximately half of all included participants had a late chronic diagnosis (51%) (Table S3), consistent with percentage of late diagnoses observed globally (19, 27, 28). Median CD4+ T cell counts in non-late diagnosis participants increased from a median of 510 cells/mm^3^ (IQR 420-630) pre-ART to 810 cells/mm^3^ (IQR 640-1120) at week 52, while in late diagnosis this increased from 215 cells/mm^3^ (IQR 90-260) to 370 cells/mm^3^ (IQR 200-540) (Table S5). Median time to viral suppression was not significantly longer in participants with a late diagnosis (31 days in non-late vs 48 in late diagnosis, p = 0.3) (Table S5). All but 2 participants remained on second generation integrase inhibitor-based therapy throughout year 1. Reservoir quantification assays and immunophenotyping were only performed in PBMCs from suppressed participants.

### Decrease in viral load and intact provirus but not inducible reservoir after one year of ART

To measure HIV-specific T cell responses following stimulation with HIV-specific peptide pools, the AIM assay was performed on PBMCs from 18 participants with HIV-1 subtype B at week 0 and week 52 (Fig. 1A). For antigen-specific CD69+4-1BB+ CD4+ T cells, a significant decrease was found after Nef (p = 0.026) and Pol (p = 0.01) stimulation (Fig. 1B, S1D), as well as in the sum of all HIV-specific responses (p = 0.043) (Fig. 1C). No significant changes in frequencies of antigen-specific OX40+CD69+CD4+ T cells were detected between week 0 and week 52 (Fig. 1D, E). 4-1BB+CD69+CD8+ T cells showed a trend to decreased responses after Pol peptide stimulation (p = 0.0506) (Fig. 1F), however both the individual peptide stimulations as well as the sum of all HIV-specific responses were not significant, possibly due to the large inter-donor variability (Fig. 1F, G).

**Figure 1:**
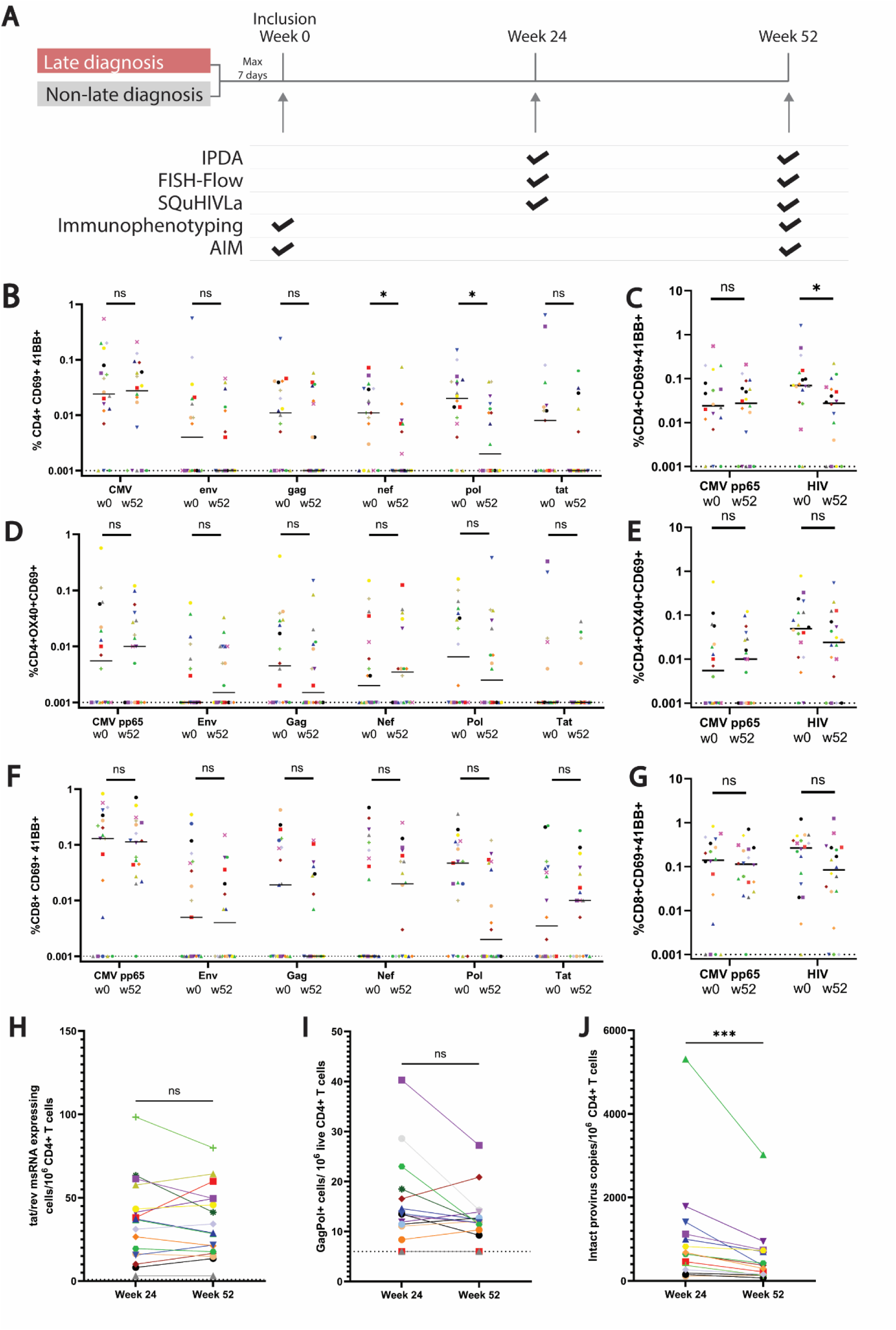
Decrease of intact viral reservoir but no significant decrease in inducible reservoir or HIV-specific T cells in the first year on ART. **A)** Schematic overview of inclusion and longitudinal sampling during the cohort duration. Arrows represent blood sampling timepoint, either by leukapheresis or phlebotomy. Participants were categorised in two groups; late diagnosis and non-late diagnosis. After diagnosis, the first sampling timepoint was maximum 7 days from initiation of ART. The ticks represent reservoir quantitation or immune characterization techniques performed on samples of that timepoint. **B-G)** AIM assay (n = 18), percentage T cells after peptide stimulation from different timepoints (week 0 and week 52 after ART initiation), percentages are normalized to DMSO-treated control. . **B)** percentage of CD4+ CD69+ 4-1BB+ T cells. Significance shown as * p<0.05. **C)** sum of CD4+CD69+4-1BB+ T cell percentages of all HIV-specific peptide stimulations. Significance shown as * p<0.05. **D)** percentage of CD4+ OX40+ CD69+ T cells **E)** sum of CD4+OX40+CD69+ T cell percentages of all HIV-specific peptide stimulations **F)** percentage of CD8+ CD69+ 4-1BB+ T cells **G)** sum of CD8+ CD69+4-1BB+ T cell percentages of all HIV-specific peptide stimulations B-G: Dashed line indicates threshold of detection. **H)** HIV-1 inducible reservoir of MS HIV-1 RNA expressing cells/10^6^ live CD4+ T cells after 12 hours of PMA+ionomycin stimulation detected by SQuHIVLa (n=17, p=0.6956) **I)** HIV-1 inducible reservoir after 18 hours of PMA+ionomycin stimulation detected by FISH-Flow (n=16) showing GagPol RNA positive cells per 10^6^ live CD4+ T cells (p=0.1272) **J)** Intact HIV-1 pro-virus detected by IPDA represented as intact HIV DNA copies/10^6^ CD4+ T cells at week 24 and week 52 after ART initiation (n=15) (p= 0.0001). Significance was calculated using a paired T-test (data normally distributed) or a Wilcoxon matched-pairs signed rank test. Normal distribution was determined with Shapiro-Wilk test. Color and icons are consistent for each participant between datasets. Due to sample availability and subtype specificity not all assays are performed for all participants.

Since ART, as expected, suppressed viral replication in these participants (Fig. S2A), we next assessed the dynamics of the (inducible) reservoir in our cohort at weeks 24 and 52 after ART initiation using SQuHIVLa and FISH-Flow as well as IPDA to quantify the intact reservoir (Fig. 1A). With SQuHIVLa, multiple spliced tat/rev RNA positive cells are detected through isothermal LAMP amplification and a maximum likelihood estimation (40). Overall, no significant decrease in the inducible HIV-1 reservoir was detected by SQuHIVLa in the first 6 months of ART-suppression, with a median of 36.78 and 28.88 tat/rev msRNA- expressing cells/10^6^ CD4+ T cells in week 24 and 52, respectively (p=0.6956) (Fig. 1H). With FISH-Flow, GagPol introns and p24 are simultaneously detected in CD4+ T cells (Fig. S1C, E) in a flow cytometry-based assay (39). FISH-Flow confirmed the absence of a significant decrease in the inducible reservoir between week 24 and week 52 (median: 13.38 and 11.88 GagPol+ cells/10^6^ CD4+ T cells, respectively) (p=0.1272) (Fig. 1I). Next, IPDA was used to detect intact proviral DNA, with digital PCR-based detection of the Psi and Env regions of the HIV genome. While the inducible reservoir did not decrease significantly, the intact proviral reservoir showed a signficant decrease between weeks 24 and 52 after ART initiation, with a median of 647 and 362 copies intact proviral DNA/10^6^ CD4+ T-cells, respectively (p= 0.0001) (Fig. 1J).

### Significant reduction of activation/exhaustion in T cell populations after one year of ART

Initial screening of clinical parameters in chronically diagnosed PWH after one year on ART showed a significant increase in CD4+ T cell count and decrease in viral load, as would be expected from ART-mediated recovery (Fig. S2A, B). To investigate immune reconstitution within the first year of ART in this chronic population in more depth, we analyzed the major blood immune cell types from 15 participants at ART initiation (week 0) and after one year on ART (week 52) using a 45-color spectral flow cytometry panel (35). To determine the cell populations, an unbiased clustering approach was chosen, allowing the detection of previously undescribed or rare cell types. For PBMCs, 24 different clusters were distinguished and annotated (Fig. S2C-K, S4, S6, Table S6). To further investigate the effect of ART specifically on T cell recovery, CD3+ cells were subclustered into 19 clusters, including exhausted (TIGIT+PD1+CD38+HLADR+) and non-exhausted CD8+ TEM clusters (Fig. 2A, S3, S5).

**Figure 2:**
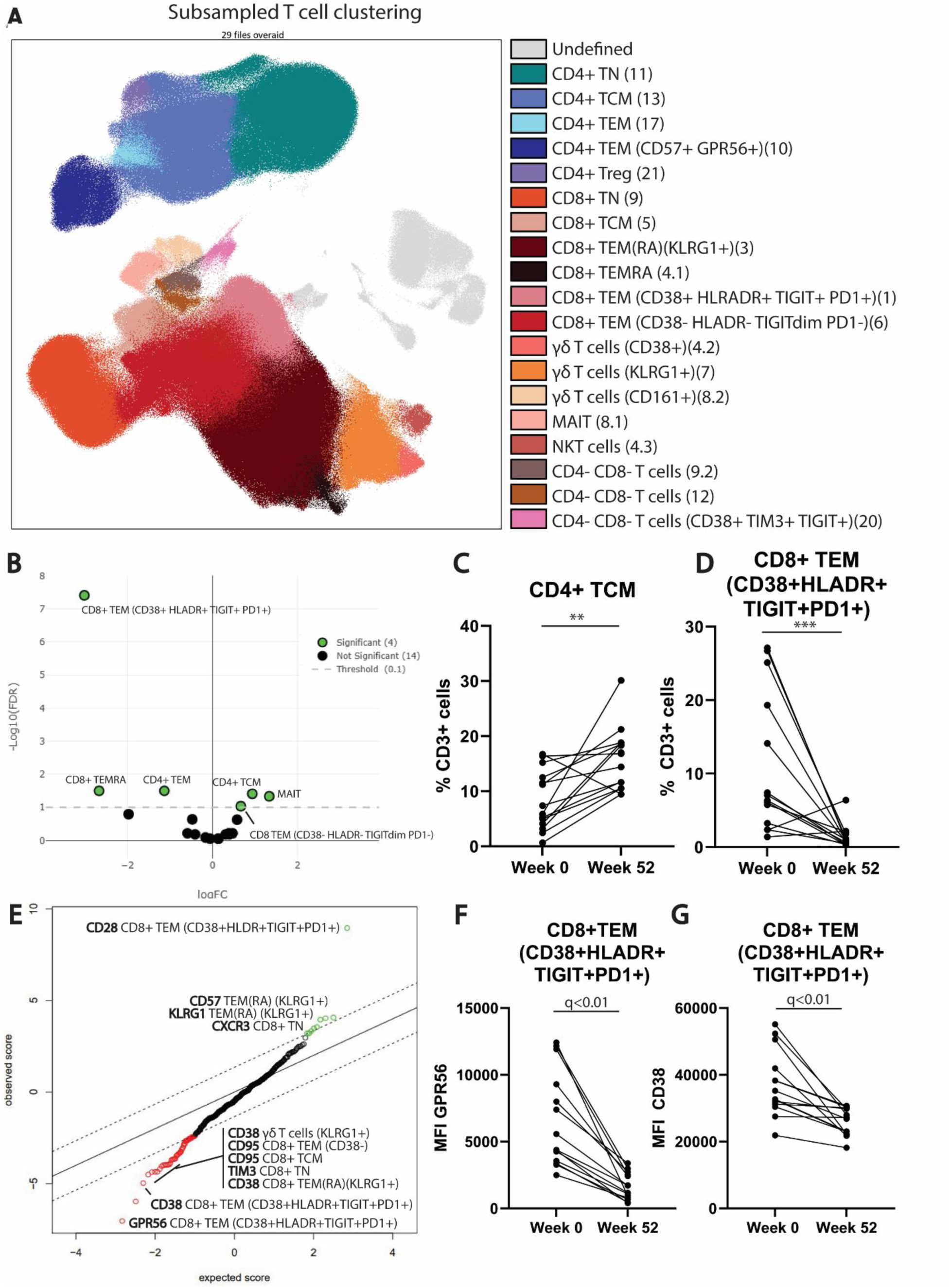
Difference in population abundance and marker expression between week 0 and week 52 within CD3+ subsampled populations. **A**) Population clustering through FlowSOM clustering algorithm (xdim; 12, ydim:12, Distance: Eucledian, clusters: 21) in high-dimensional 45-color flow cytometry data. Dimensionality reduction and visualization by UMAP (Neighbors: 80, Minimum distance: 0.7, Distance: Euclidean, Epochs: 251), all samples concatenated. **B**) Volcanoplot of differences in cluster abundance between week 0 and week 52, green color indicates significance (EdgeR, FDR: 0.1) positive represents higher abundance at week 52 than week 0 after ART initiation, negative lower. **C**) Percentage of subsampled CD3+ cells classified as CD4+ central memory T cell cluster in week 0 (n=14) and week 52 (n=15)(** FDR < 0.05). **D**) percentage of subsampled CD3+ cells that were exhausted CD8+ effector memory T cells (CD38+ HLADR+ TIGIT+ PD1+ ) at week 0 (n=14) and week 52 (n=15)(*** FDR < 0.001). **E**) SAM analysis of differential activation/exhaustion marker expression on CD3+ subsampled clusters, with green indicating higher expression at week 52 and red lower expression at week 52 compared to week 0 (q-value <0.01, number of permutations: 500). **F**) MFI of CD38 on CD8+ effector memory T cell cluster (CD38+HLADR+) at week 0 and week 52 (n=14, q-value <0.01). **G**) MFI of PD1 on CD4+ effector memory T cell cluster at week 0 and week 52 (n=14, q-value <0.01).

**Figure 3.**
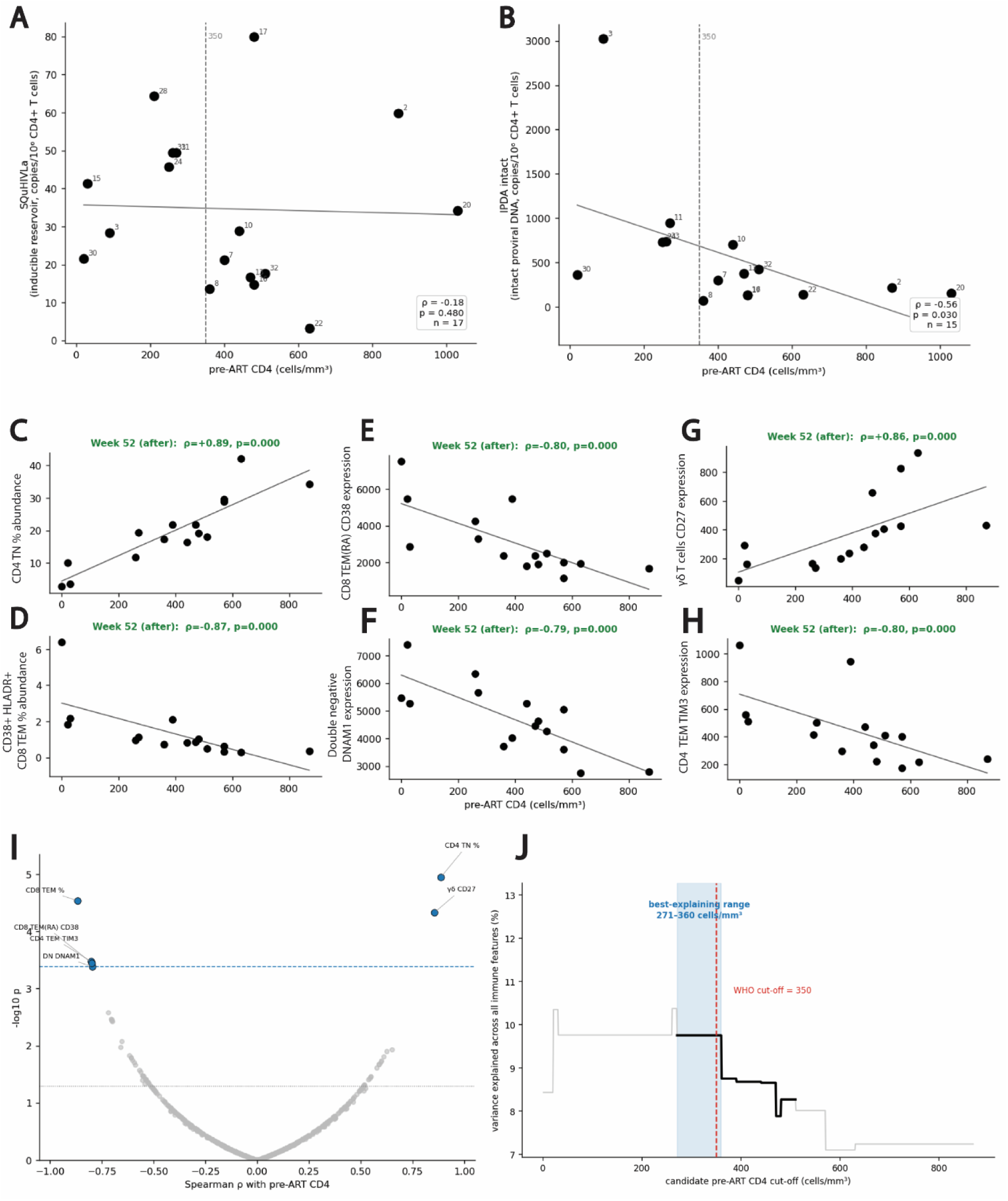
Correlations with pre-ART CD4. **A**) Correlation of pre-ART CD4 with HIV-1 inducible reservoir of MS HIV-1 RNA expressing cells/10^6^ live CD4+ T cells after 12 hours of PMA+ionomycin stimulation detected by SQuHIVLa at week 52 after ART initiation (n = 17, p = 0.480), **B)** Correlation of pre-ART CD4 with intact HIV DNA copies/10^6^ CD4+ T cells at week 52 after ART initiation (n = 15, p = 0.03) **C-H)** Individual correlations of significant features, MFI on Y-axis, pre-ART CD4 (cells/mm^3^) on X-axis. **I)** Correlation of pre-ART CD4 with all immune features (MFI and abundance) tested at the same time with one BH-FDR correction across all features (n = 15, FDR 0.05). **J**) Data-driven optimal split of the cohort into two groups with pre-ART CD4 cut-off, based on variance explained across all immune features.

Next, we examined the differences in abundance of PBMC clusters at weeks 0 and 52 (Fig. S6A-C). As expected from previous reports, we observed a significant increase in CD4+ central memory T cells (TCM), mucosal-associated invariant T cells (MAIT), plasmacytoid dendritic cells (pDC), conventional dendritic cells (cDC) and naïve B cells, while the abundance of CD8+ effector memory T cells (TEM) decreased at week 52 (Fig. S6C-E)(43–45). T cell clustering confirmed the increase of MAIT and CD4 TCM (Fig 2B, C). Interestingly, within T cell clustering, a specific exhausted CD8+ TEM (CD38+ HLADR+ TIGIT+ PD1+) population decreased in abundance at week 52 after ART initiation, whereas non-exhausted CD8+ TEM were increased in abundance, demonstrating that the decrease in CD8+ TEM is mostly driven by the exhausted CD8+ TEM population that was not identified with full PBMC clustering (Fig. 2B, D). Therefore, while the conclusions remain generally consistent between T cell and PBMC analysis, subclustering allows for a more in-depth analysis of cluster phenotypes.

In addition to cell subset abundances, determining marker expression within each cluster reveals more detailed information about immune reconstitution after one year on ART. Expression of several activation and exhaustion markers decreased on various T cell subsets after one year of ART. For instance, in PBMC clustering, expression of activation/exhaustion markers CD38, GPR56, HLA-DR and PD1 on CD8+ TEM was decreased (Fig. S6F). T cell subsampling analysis confirmed the decrease in CD38 and GPR56 expression on exhausted CD8+ TEM. Moreover, the increase in CXCR3 and decrease in TIM3 expression previously determined on CD8+ TN in the PBMC subsampling were confirmed (Fig. 2E-G).

Notably, most of the observed population abundance changes have been previously reported in HIV-1 recovery after ART, thereby reaffirming the validity and robustness of this technology, while providing more in-depth information about dynamics of activation and exhaustion profiles during ART (43–48).

### Variability in immune and reservoir components correlates with pre-ART CD4

We hypothesized that the large inter-individual variability of HIV-1 reservoir, immune cell abundance and immune marker expression at week 52 was due to heterogeneity within the group of PWH diagnosed at a chronic stage. To examine if this heterogeneity can be correlated with a known and regularly tested clinical parameter, correlations with pre-ART CD4+ T cell count were tested. The amount of inducible HIV-1 msRNA expressing cells at week 52 did not correlate with pre-ART CD4+ T cell count (Fig. 3A), however intact HIV-1 proviral DNA (IPDA) did correlate significantly with pre-ART CD4+ T cell count (p = 0.03)(Fig. 3B).

We then tested all markers and cell populations (522 features) from the immunophenotyping dataset within four T cell compartments (αβCD4+, αβ CD8+, CD4- CD8- and non-αβ T cells) for a correlation with CD4+ T cell count pre-ART and inducible HIV-1 reservoir (SQuHIVLa) (Fig. 3I, S7A). No statistically significant correlations with SQuHIVLa were observed after adjustment for multiple testing (Fig. S7A). Pre-ART CD4+ T-cell counts correlated positively with the abundance of CD4+ TN at week 52, and negatively with exhausted CD8+ TEM abundance (Fig. 3C-D). Moreover, the expression of CD27 on γδ T cells, CD38 on TEM(RA), TIM3 on CD4+ TEM and DNAM on DN correlated significantly with pre-ART CD4+ T cell count (Fig. 3E-I). PCA was performed across all subsampled T-cell populations to determine whether individuals with high and low CD4 counts formed distinct clusters. Samples did not separate based on pre-ART CD4 count, possibly due to other variables in the dataset such as sex and age (Fig. S7B).

Since inclusion and exclusion criteria require a pre-set cutoff, we set out to determine a division of subgroups to best describe the variability within the population with a chronic HIV-1 diagnosis. Therefore, we modelled the best division within the variability of all immunophenotyping data, which indicated that the division that explained the variability the best would be between 271 and 360 CD4+ T cells/mm^3^ (Fig. 3J). This also corresponds to the WHO definition of late (<350 CD4+ T cells/mm^3^ or an AIDS-defining illness) and non-late diagnosis subgroups. Using these WHO criteria, the cohort was divided into participants with a late diagnosis and a non-late diagnosis.

### Higher inducible reservoir and abundance of exhausted CD8+ T cells with late diagnosis after one year ART

To further explore the difference between the two subgroups of chronic diagnosis, we first compared the inducible reservoir between late and non-late chronic diagnosis. Interestingly, we observed a significantly higher inducible msRNA expressing reservoir in the late diagnosis subgroup both at week 24 (p= 0.0044) (Fig. 4A) as well as week 52 (p=0.012)(Fig. 4B) as determined by SQuHIVLa. The intact proviral reservoir was also significantly higher in the late diagnosis subgroup at week 24 (p = 0.012) (Fig. S7C) with a higher trend also observed at week 52, though this was not significant, possibly due to the low participant number (p = 0.066) (Fig. S7D). Analysis of inducible cell-associated GagPol transcripts by FISH-Flow did not show a significant difference between the two subgroups (Fig. S7E-F).

**Figure 4:**
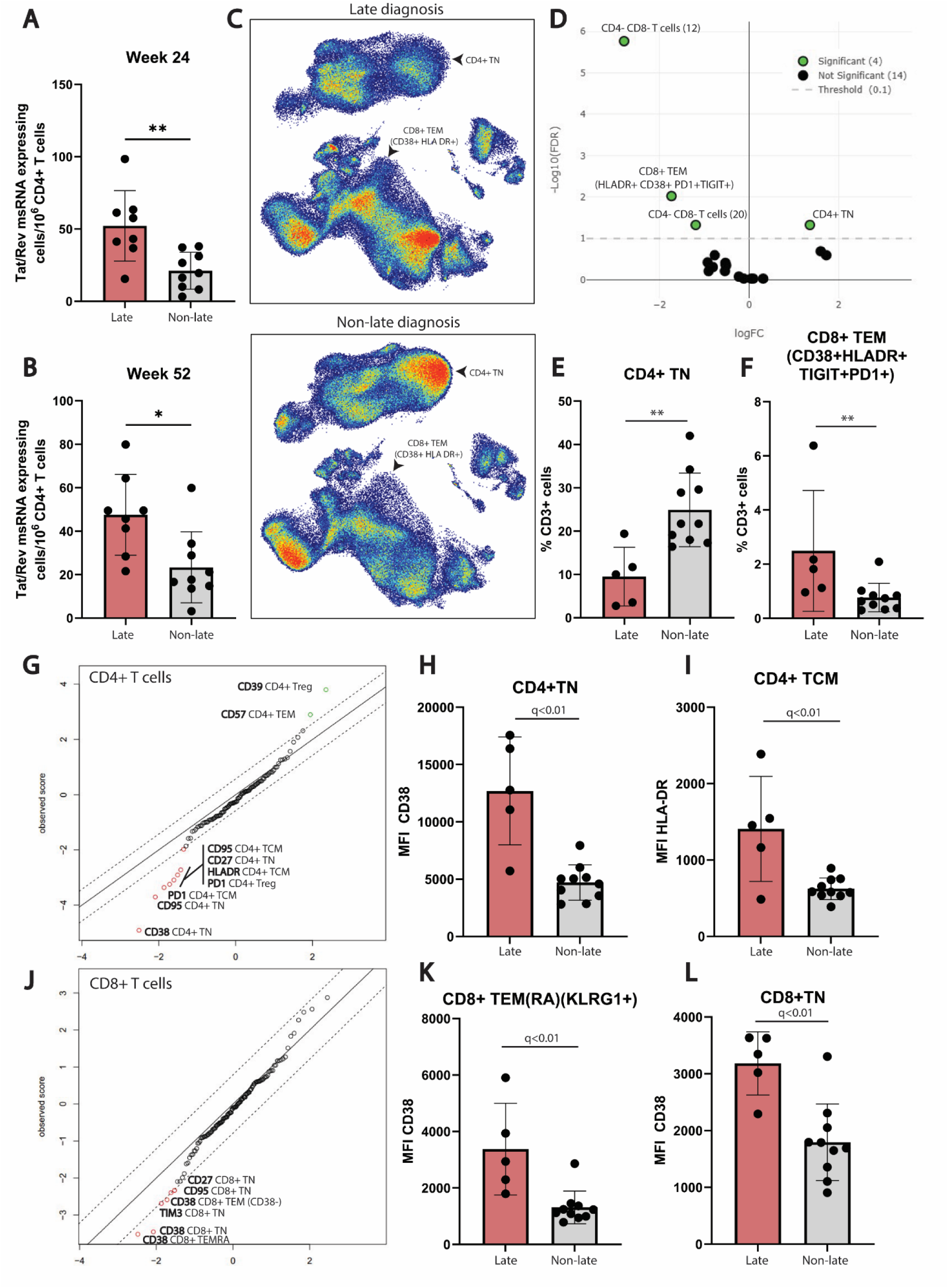
Reservoir inducibility and immune phenotypic differences between the late and non-late diagnosis groups. **A-B)** Inducible reservoir at week 24 in late (n=8) and non-late (n=9) diagnosis group shown as Tat/rev msRNA expressing cells/10^6^ CD4+ T cells measured by SQuHIVLa after 12 hours of PMA/ionomycin stimulation (** p<0.01, * p<0.05) at **A)** week 24 **B)** week 52 **C)** UMAP depiction of CD3+ subsampled cells in concatenated samples from late (n=5) and non-late (representative n=5) group. Differences in cluster abundances are shown with black arrows. **D)** Volcano plot of significant differences in cluster abundance between late (n=5) and non-late (n=10) diagnosis groups. Green dots are significant (FDR: 0.1) positive represents higher abundance in the non-late than in the late group, negative represents higher abundance in the late than in the non-late group. **E)** Percentage of CD3+ cells classified as CD4+ naïve T cell cluster in the late (n=5) and non-late (n=10) diagnosis groups (** FDR< 0.05). **F)** Percentage of CD3+ cells classified as exhausted CD8+ effector memory T cell cluster (CD38+HLADR+TIGIT+PD1+) in the late (n=5) and non-late (n=10) diagnosis groups (** FDR< 0.05). **G)** SAM analysis of activation/exhaustion markers on CD4+ T cell clusters between late (n=5) and non-late (n=10) diagnosis, green values represent higher marker expression in the non-late diagnosis group, red represent lower expression in the non-late group (q-value <0.01). **H)** MFI of CD38 expression on CD4+ naïve T cell cluster in late (n=5) and non-late (n=10) diagnosis group (q-value < 0.01). **I)** MFI of HLA-DR expression on CD4+ central memory T cell cluster in late (n=5) and non-late (n=10) diagnosis group (q-value < 0.01). **J)** SAM analysis of activation/exhaustion markers on CD8+ T cell clusters between late (n=5) and non-late (n=10) diagnosis group, green values represent higher marker expression in the non-late diagnosis group, red represent lower expression in the non-late group (q-value <0.01). **K)** MFI of CD38 expression on CD8+ TEMRA cluster in late (n=5) and non-late (n=10) diagnosis groups (q-value < 0.01). **L)** MFI of CD38 expression on CD8+ naïve T cell cluster in late (n=5) and non-late (n=10) diagnosis groups (q-value < 0.01). All statistical analysis and multiple-testing correction was performed in OMIQ-integrated EdgeR.

To investigate if the differences observed in the reservoir size and inducibility match differences in the immune compartment, we compared cluster abundances between the late and non-late diagnosis subgroups after one year on ART. When clustering all PBMCs, abundance of CD4+ TN was significantly lower in the subgroup with a late diagnosis (Fig. S7G). As expected, after T cell sub clustering, CD4+ TN abundance was also found to be significantly lower in the subgroup with a late diagnosis (Fig. 4C,D, E). Within the subgroup with a late diagnosis, we also observed a significantly higher abundance of exhausted TIGIT+ PD1+ CD38+ HLADR+ CD8+ TEM (Fig. 4C, D, F) at one year of ART suppression. Interestingly, the non-exhausted TIGIT-PD1-CD38-HLADR- CD8+ TEM showed no significant difference between the late and non-late diagnosis subgroups (Fig. 4D).

Next, we determined whether the expression of additional exhaustion and activation markers was different between the late and non-late diagnosis subgroups a year after ART initiation. CD95 (Fas) expression on CD8+ TN, CD4+ TN, and CD4+ TCM was higher in PWH with late diagnosis (Fig. 4G, J). Higher CD95 expression on CD8+ TN coincided with a significantly higher inhibitory marker TIM3 (Fig. 4J) and activation marker CD38 (Fig. 4J, L) expression when compared to the non-late diagnosis subgroup. Additionally, the late diagnosis subgroup exhibited higher expression of CD38 on CD8+ TEMRA, non-exhausted CD8+ TEM and CD4+ naïve T cell cluster (Fig. 4G, H, J, K). Lastly, CD4+ regulatory T cells in the non-late diagnosis subgroup showed higher expression of CD39 (Fig. 4G). Antigen-specific T cell responses did not show significant differences between the late and non-late diagnosis subgroups (Fig. S7H-M). Overall, both cluster abundance as well as marker expression showed more exhaustion of T cell populations in the late than in the non-late diagnosis subgroups after one year on ART.

## Discussion

In this study, we investigate reservoir and immune compartment dynamics in non-acutely diagnosed PWH during their first year on ART. While the clinical distinction is often made between acute (< Fiebig VI) and chronic (from Fiebig VI) diagnosis, there is great heterogeneity within chronic diagnoses. To obtain more insight into this heterogeneity, we stratified participants into late (CD4+ T cell count < 350 cells/mm^3^ or an AIDS-defining illness at diagnosis) and non-late diagnosis and treatment. After one year on ART, the inducible reservoir measured by SQuHIVLa as well as T cell exhaustion measured with in-depth immunophenotyping were higher in PWH with a late diagnosis (Fig. 4). Overall, our results demonstrate the importance of timely diagnosis and treatment of PWH, with possible implications for trials testing interventions.

Consistent with previous data, we found a decrease in HIV-specific T cells between week 0 and week 52, as would be expected following a decrease in antigen exposure (49). Our findings show no decrease in the inducible reservoir in both gag-pol unspliced RNA and Tat/Rev multiple spliced RNA between week 24 and week 52, consistent with previous observations that the reservoir decays slower in PWH treated at a chronic stage (Fig. 1H- I)(16, 50). The intact HIV-DNA (IPDA) did indicate a decrease between weeks 24 and 52, in line with other studies that demonstrated a decrease of the intact proviral reservoir within the first year on ART (Fig. 1J)(25, 51, 52). The two techniques detect different reservoir compartments, with IPDA detecting intact DNA whereas SQuHIVLa detects inducible multiple spliced RNA and requires a 12-hour stimulation, which could contribute to the difference in findings. Our observation that the inducible reservoir remained unchanged between 24 weeks and 52 weeks after ART initiation, while the amount of intact HIV DNA decreased during the same period might suggest that the IPDA assay detected some productively infected cells that were eliminated within the first 12 months. However, our data provide only indirect evidence, as we did not perform assays to detect spontaneous provirus expression in these cells. Another possible explanation is that the first decay phase of the inducible reservoir is shorter than that of the HIV DNA reservoir, and was already completed before week 24, as was recently shown with inducible HIV-protein by Puertas et al (53). This observation warrants further investigation as accelerating the elimination not only of the intact but also of the inducible reservoirs may provide clinical benefits as well as open new cure opportunities.

Deep immunophenotyping showed several changes after one year of ART. First, the abundance of exhausted CD38+HLA-DR+TIGIT+PD1+CD8+ TEM decreased (Fig. 2B). This phenotype for CD8+ TEM was previously reported to be present at higher abundance in PWH at later stages of disease (54). CD38 and HLA-DR have been associated with progression to AIDS (55, 56) and shorter survival in PWH with late-stage AIDS (57) as well as in other viral infections (58, 59). In accordance with literature, CD4+ TCM, MAIT cells, pDCs, cDCs, CD8+ TCM and naïve B cells increased in abundance, whereas marginal zone-like B cells decreased in abundance (43, 44, 46, 47, 60–62) (Fig. S6C). The fact that many of our findings have previously been reported using conventional flow cytometry and manual gating further supports the robustness of the multiparameter spectral flow cytometry as well as the unsupervised clustering.

We found that the intact and inducible reservoir in the late diagnosis group is higher than in the non-late diagnosis group both at week 24 and week 52 after ART initiation (Fig. 4, Fig. S7). Previous reports observed that a higher CD4+ T cell count at time of diagnosis correlated with lower HIV-1 DNA after suppression, in line with our observations of a lower reservoir in PWH with a non-late diagnosis (63). In PWH with a chronic diagnosis, higher CD4+ T cell nadir was associated with faster intact proviral DNA decay (64) and a higher likelihood of having HIV-DNA <100 copies/10^6^ PBMCs (65). However, all of the aforementioned studies quantified HIV-1 DNA, without directly reflecting the transcriptionally-competent reservoir. In contrast, in our study, we examined the difference between late and non-late populations through characterization of the viral reservoir in different molecular compartments. This included SQuHIVLa, which quantifies the inducible reservoir expressing multiple spliced RNA, with multiple spliced RNA suggested to be a good predictor for viral rebound after treatment interruption (40, 66).

Importantly, using our high dimensional multi-parameter spectral flow cytometry approach, we could point out several interesting differences within the immune compartment between the late and non-late diagnosis subgroups. Most notably, the population of exhausted CD8+ TEM (CD38+HLADR+PD1+TIGIT+) was significantly higher in abundance in the late diagnosis subgroup (Fig. 4). Moreover, the abundance of CD4+ naïve T cells remained higher in the non-late diagnosis subgroup, even after one year of ART. Crucially, these findings demonstrate that although PWH with a late diagnosis show recovery on ART in general clinical parameters (CD4+ T cell count, viral load), immune exhaustion remains higher and naïve CD4+ T abundance lower in the late diagnosis subgroup.

Beside the exhausted CD8+ TEM population, exhaustion in the late diagnosis subgroup was illustrated by higher expression of exhaustion markers on other cell populations. For instance, expression of CD38 and PD1, both previously reported exhaustion markers (67–69), was higher on CD8+ TEMRA and CD4+ TCM cells respectively from the late diagnosis subgroup. We also found CD38 to be more highly expressed on CD8+ naïve T cells and CD4+ naïve T cells from the late diagnosis subgroup one year after ART initiation (Fig. 4). While CD38 is an activation/exhaustion marker on effector cells, on naïve cells it was reported to indicate recent thymic emigration, suggesting that the late diagnosis subgroup has a higher thymic output (70). Moreover, CD4+ Tregs expressed more CD39 in the non-late diagnosis subgroup, possibly increasing their function and allowing for the reduction of chronic activation (71). Overall, increased exhaustion marker expression on several populations indicates a higher state of exhaustion in the late diagnosis subgroup. Interestingly, the exhausted CD8+TEM population correlated significantly with the CD4+ T cell count pre-ART, providing further evidence for the large impact of time to diagnosis on activation and exhaustion after a year on therapy.

CD95 (Fas) expression was higher on CD4+ TCM, CD4+ TN and CD8+ TN cells of PWH from the late diagnosis subgroup (Fig. 4). This could be due to higher residual expression of HIV proteins in PWH from the late diagnosis subgroup, leading to T cell activation and CD95 expression. Although mostly known for its induction of apoptosis, CD95 can also be an activation marker (72). Typically, naïve T cells express low to no CD95, as CD95 expression is induced upon T cell activation. However, CD95 expression on naïve CD4+ T cells has been reported to increase with HIV disease progression (73). Additionally, CD95 stimulation on naïve T cells was described to inhibit proliferation and activation of naïve T cells (74). Without functional studies, the specific role CD95 plays in this context remains unknown.

A limitation of this study is that while the week 0 to week 52 comparison is longitudinal, the late- versus non-late analysis is non-paired, making it more vulnerable to age-, subtype- and sex-based biases. We have performed several analyses to mitigate these biases, however without perfect matching, unfortunately impossible due to limited sample availability, it cannot be excluded that confounding factors such as sex and age may have influenced this analysis. Moreover, while our cohort is representative of people living with HIV in the Netherlands, findings may not be generalizable to other settings.

While long-term decay of the HIV-1 reservoir has been extensively studied and shown to occur at a very slow rate, much less is known about the dynamics within the first year of ART (75). This is a critical gap, especially as an increasing number of HIV cure strategies are being designed to intervene at or shortly after ART initiation (76). As new technologies such as single-cell multi-omics, spatial transcriptomics, and improved latency assays continue to evolve, this ongoing cohort will serve as a valuable platform for applying and validating these tools. In particular, understanding the immunological and virological trajectories of individuals initiating ART at different stages, such as late versus non-late, will be critical for personalizing cure strategies and ensuring equitable progress toward a functional cure. Importantly, in the design of reservoir quantitation studies and HIV-1 cure trials it is essential to consider time of diagnosis, especially also within chronic diagnoses, as starting points and expected dynamics of both the persistent reservoir and immune compartment are different between the late and non-late diagnosis subgroups.

## Contributors

Conceptualization, K.H., L.V., Y.M, C.R., T.Ma, Methodology, L.V., K.H., T.H., J.O., R.C., A.G., Investigation, L.V., K.H, T.H., J.O., Data Curation and Formal Analysis; L.V., K.H., M.B., Y.M, C.R., T.Ma, Data Visualization; K.H, L.V., M.B., Y.M, C.R., T.Ma, Project Administration; K.H., L.V., Y.M, C.R., T.Ma, Supervision; Y.M, C.R., T.Ma, Material and Funding Acquisition; K.H., A.G., Y.M, C.R, T.M. Writing – Original draft; L.V., K.H., Writing – Review and Editing; L.V., K.H., T.H., J.O., R.C., M.B., A.G., C.L., R.G., R.P., D.V., J.K., P.K., T.Me., S.R., Y.M., C.R., T.Ma.

## Acknowledgements

We extend our deepest gratitude to all the people living with HIV (PLWH) who participated in our study. We also wish to express our sincere thanks to Erik Gausvik, Marianne van Wingerden, and Safana Safanatunnajah for their management and inclusion of the cohort. We want to express our appreciation to Marjan van Meurs and Danique Laport for assistance with multi-color spectral flow cytometry. Our thoughts go to the late prof. Charles Boucher who was a driving force for this study.

## Data sharing statement

The presented and additional data from this study can be shared on request. Please contact the corresponding author for further information

