## Supplementary for "Larger inducible reservoir and higher abundance of exhausted CD8+ T cells in people treated after late versus non-late diagnosis of chronic HIV one year after ART initiation"

### Supplementary material

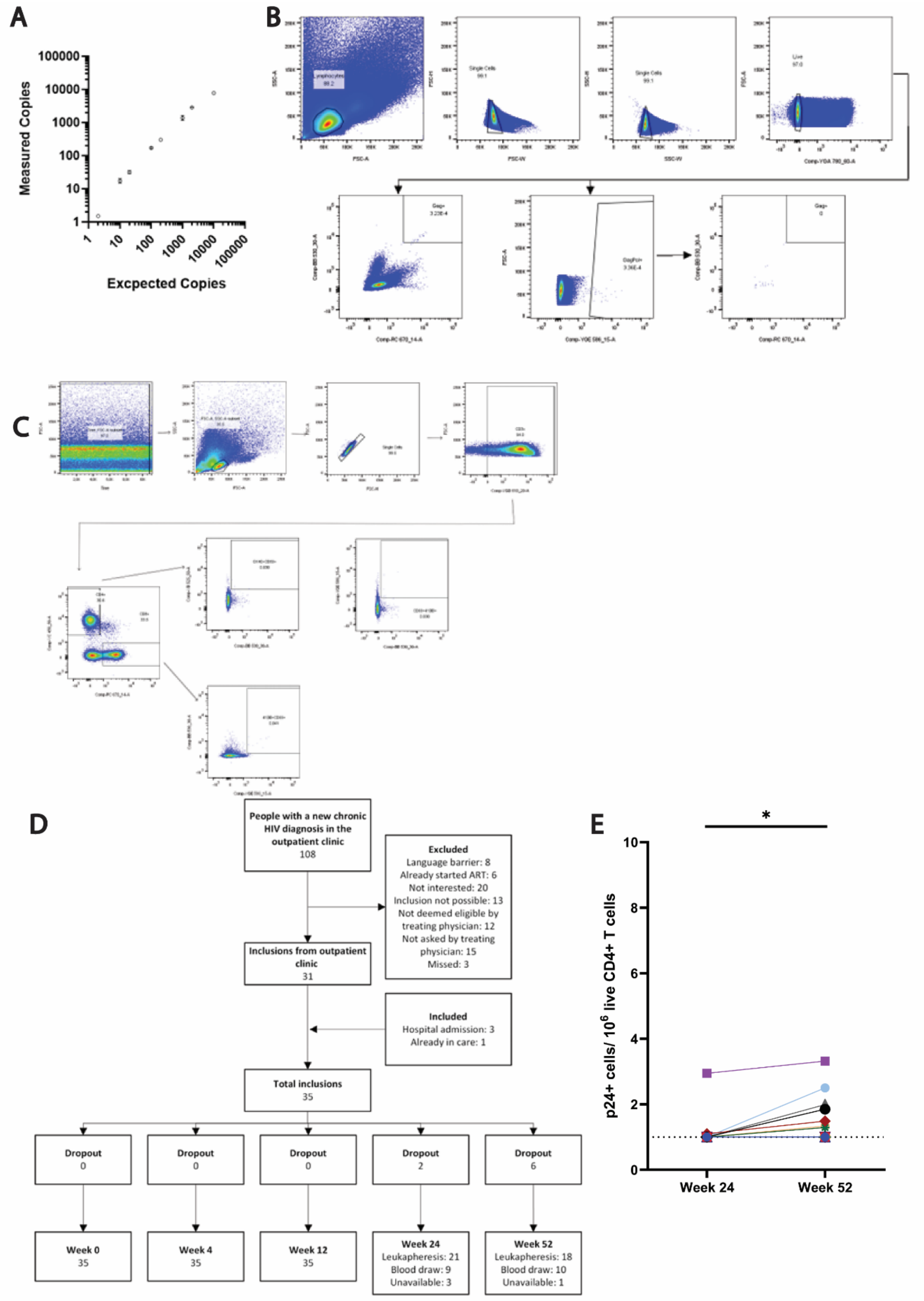

**Fig S1 A)** Validation of novel IPDA primers and probes using the pNL4.3 plasmid . Data are shown as the mean ± standard deviation of two serial dilutions, with up to two technical replicates per dilution. **B)** Gating strategy Fluorescent in situ hybridization flow cytometry (FISH-Flow) **C)** Gating strategy Activation Induced Marker Assay. **D)** Flowchart of participant inclusions. **E)** Inducible translationally active reservoir as measured by p24+ cells/$10^6$ live CD4+ T cells at weeks 24 and 52 ($p = 0.016$), Wilcoxon matched-pairs signed rank test.

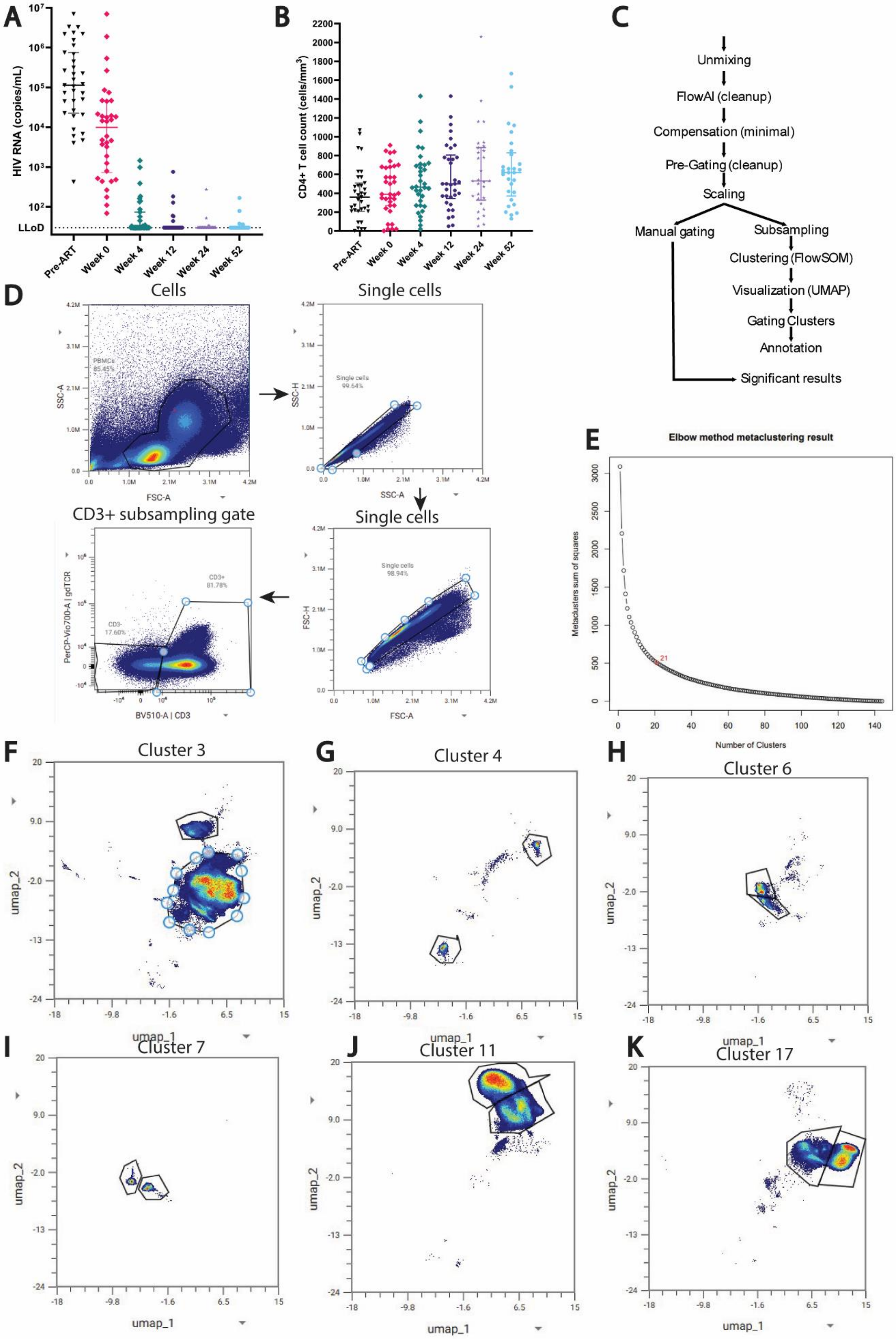
A
HIV RNA (copies/mL)
LLoD
Pre-ART
Week 0
Week 4
Week 12
Week 24
Week 52
B
CD4+ T cell count (cells/mm3)
C
Unmixing
FlowAI (cleanup)
Compensation (minimal)
Pre-Gating (cleanup)
Scaling
Manual gating
Subsampling
Clustering (FlowSOM)
Visualization (UMAP)
Gating Clusters
Annotation
Significant results
D
Cells
Single cells
CD3+ subsampling gate
Single cells
PBMCs 85.45%
Single cells 99.64%
Single cells 98.94%
CD3+ 81.78%
CD3- 17.60%
SSC-A
FSC-A
SSC-H
FSC-H
PerCP-Vio700-A | gdTCR
BV510-A | CD3
E
Elbow method metaclustering result
Metaclusters sum of squares
Number of Clusters
F
Cluster 3
G
Cluster 4
H
Cluster 6
I
Cluster 7
J
Cluster 11
K
Cluster 17
umap_1
umap_2

**Fig S2 A)** HIV RNA viral load in plasma (copies/mL) pre-ART and at several timepoints after ART initiation **B)** CD4+ T cell count (cells/mm$^3$) at baseline and specified time points after ART initiation **C)** Immunophenotyping sample processing in OMIQ **D)** Gating strategy cleanup and CD3 subsampling immunophenotyping **E)** FlowSOM elbow metaclustering result for all PBMCs **F-K)** Subclustering of **F)** cluster 3 **G)** cluster 4 **H)** cluster 6 **I)** Cluster 7 **J)** Cluster 11 **K)** Cluster 17.

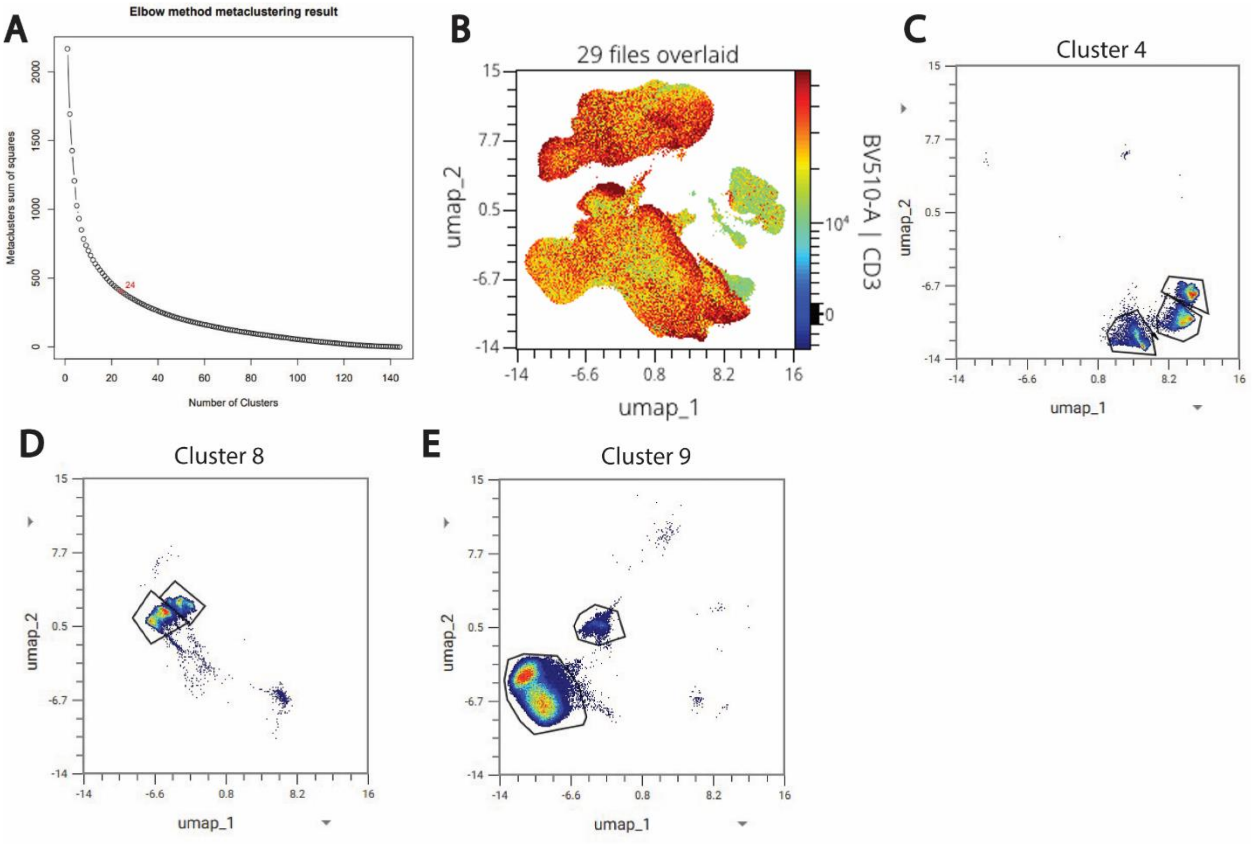


**Fig S3. A)** FlowSOM elbow metaclustering analysis for subsampled T cells **B)** Colour-continous scatterplot UMAP of subsampled T cells with CD3 expression on Z-axis **C-E)** Manual subclustering after CD3 subsampling of **C)** cluster 4 **D)** cluster 8 **E)** Cluster 9.

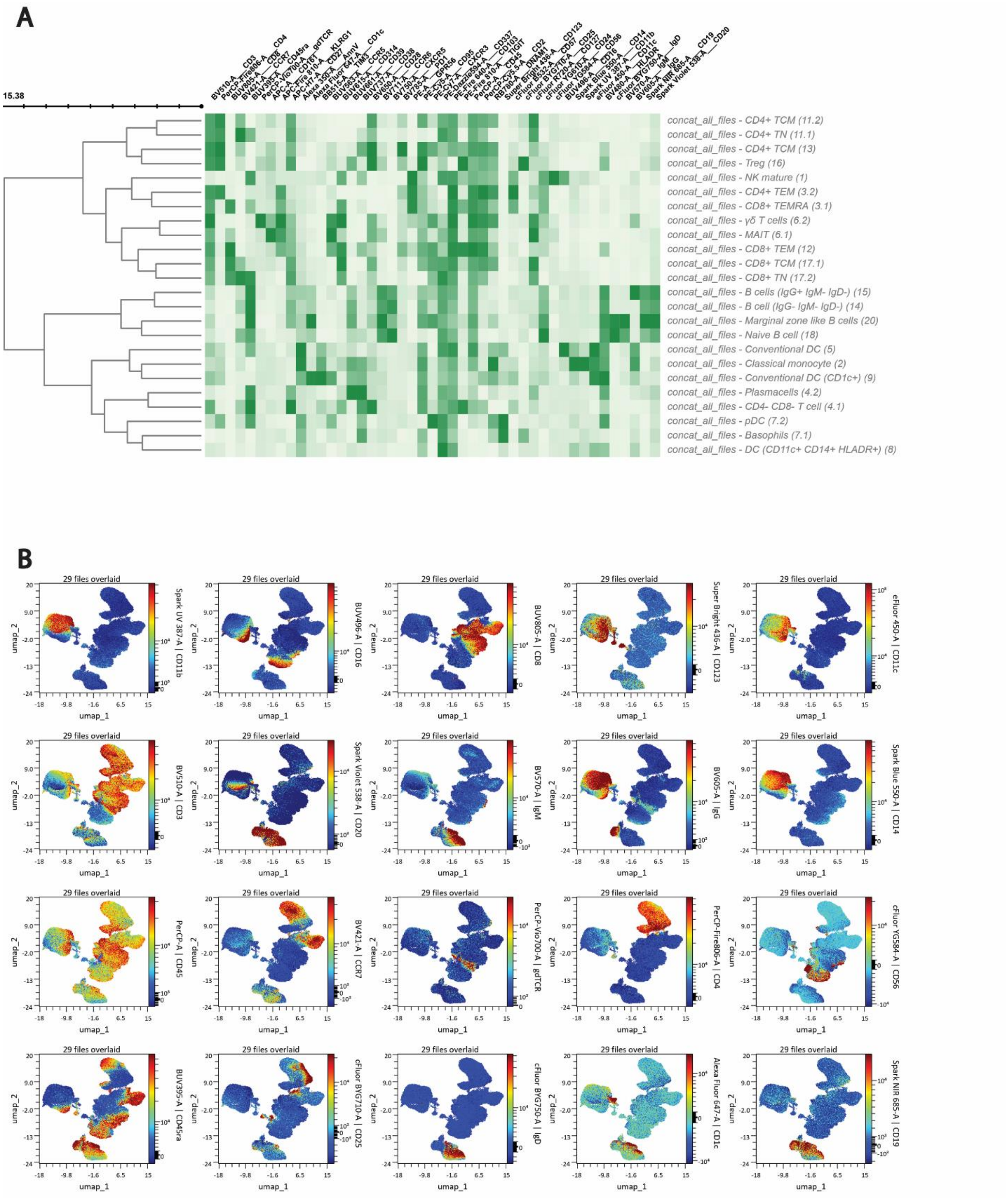

**Fig. S4. A)** Heatmap of PBMC cluster expression of each marker **B)** UMAP of PBMC clustering with expression of major lineage markers.

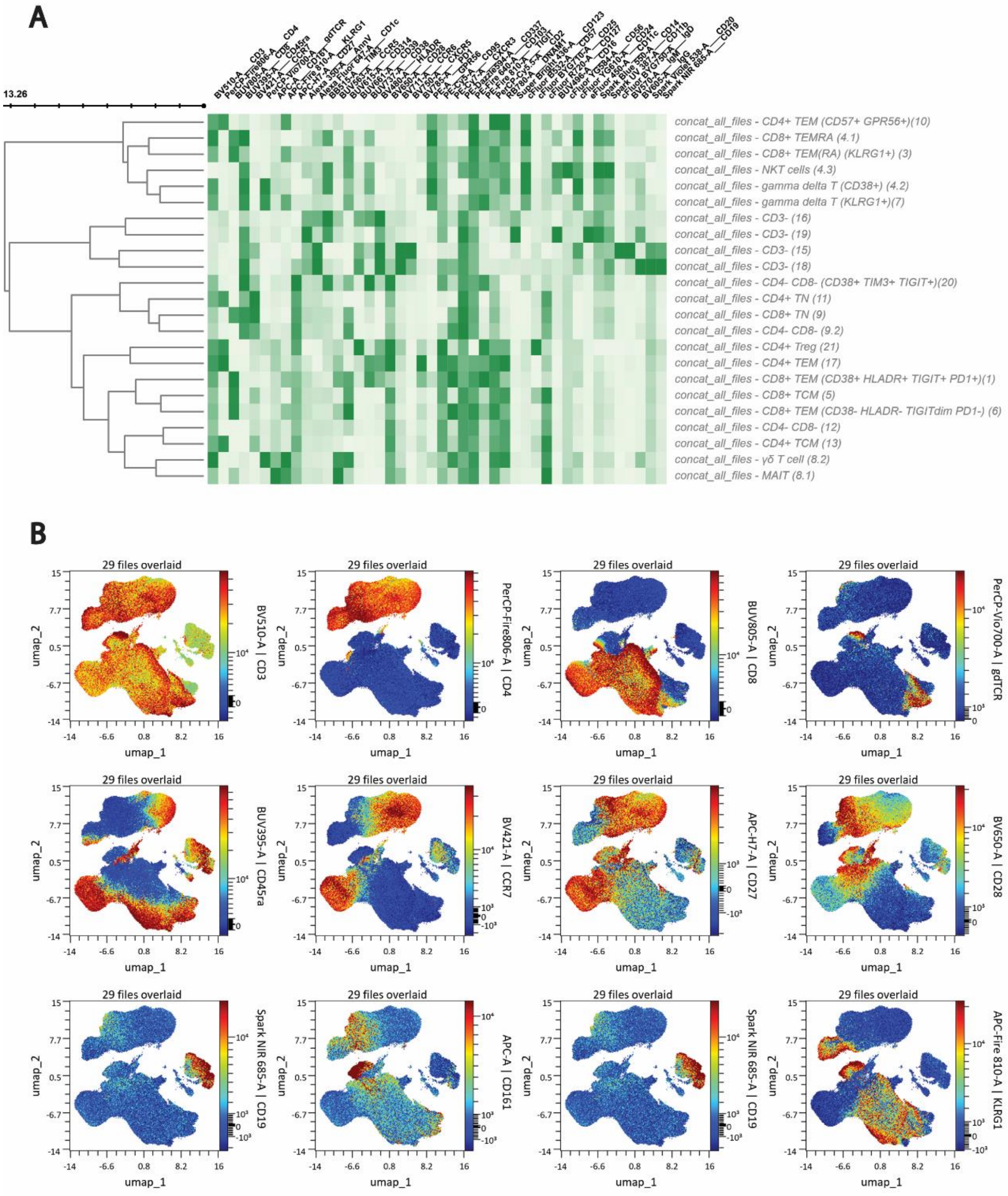


**Fig S5. A)** Heatmap of CD3 subsampled cluster expression of each marker **B)** UMAP of CD3 subsampled clustering with expression of major lineage markers.

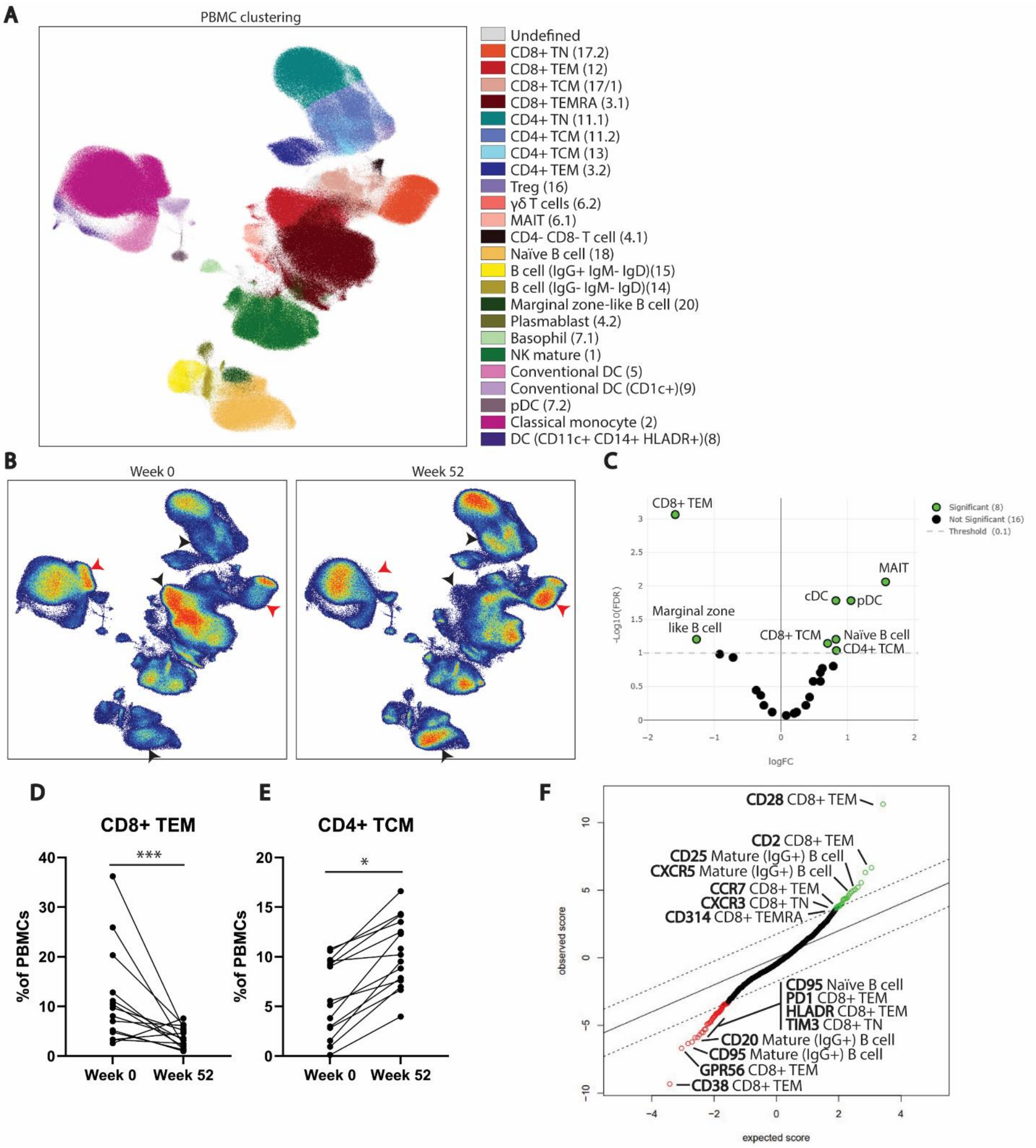


**Fig S6**. **Annotation of cell populations in PBMCs A**) Population clustering through FlowSOM clustering algorithm (xdim; 12, ydim:12, Distance: Eucledian, clusters: 20) in high-dimensional 45-color flow cytometry data. Dimensionality reduction and visualization by UMAP (Neighbors: 80, Minimum distance: 0.7, Distance: Euclidean, Epochs: 250), all samples concatenated. **B**) UMAP visualization of week 0 (n=14) and week 52 (n=14, representative), abundance differences are pointed out with black arrows, differences within the same cluster are pointed out with red arrows. **C**) Volcano plot of population abundance differences between week 0 and week 52 (EdgeR, FDR: 0.1), positive representing higher abundance at week 52 than week 0 after ART initiation, negative lower. **D**) CD8+ TEM cluster abundance in percentage of PBMCs at week 0 (n=14) and week 52 (n=15)(*** FDR < 0.001) **E**) CD4+ TCM cluster abundance in percentage of PBMCs at week 0 (n=14) and week 52 (n=15)(* FDR < 0.1) **F**) SAM analysis of differential median marker expression (excluding CD45 and Annexin V) on PBMC clusters between week 0 and week 52, green indicating higher expression at week 52 compared to week 0, red indicating lower expression at week 52 (q-value <0.01, number of permutations: 500).

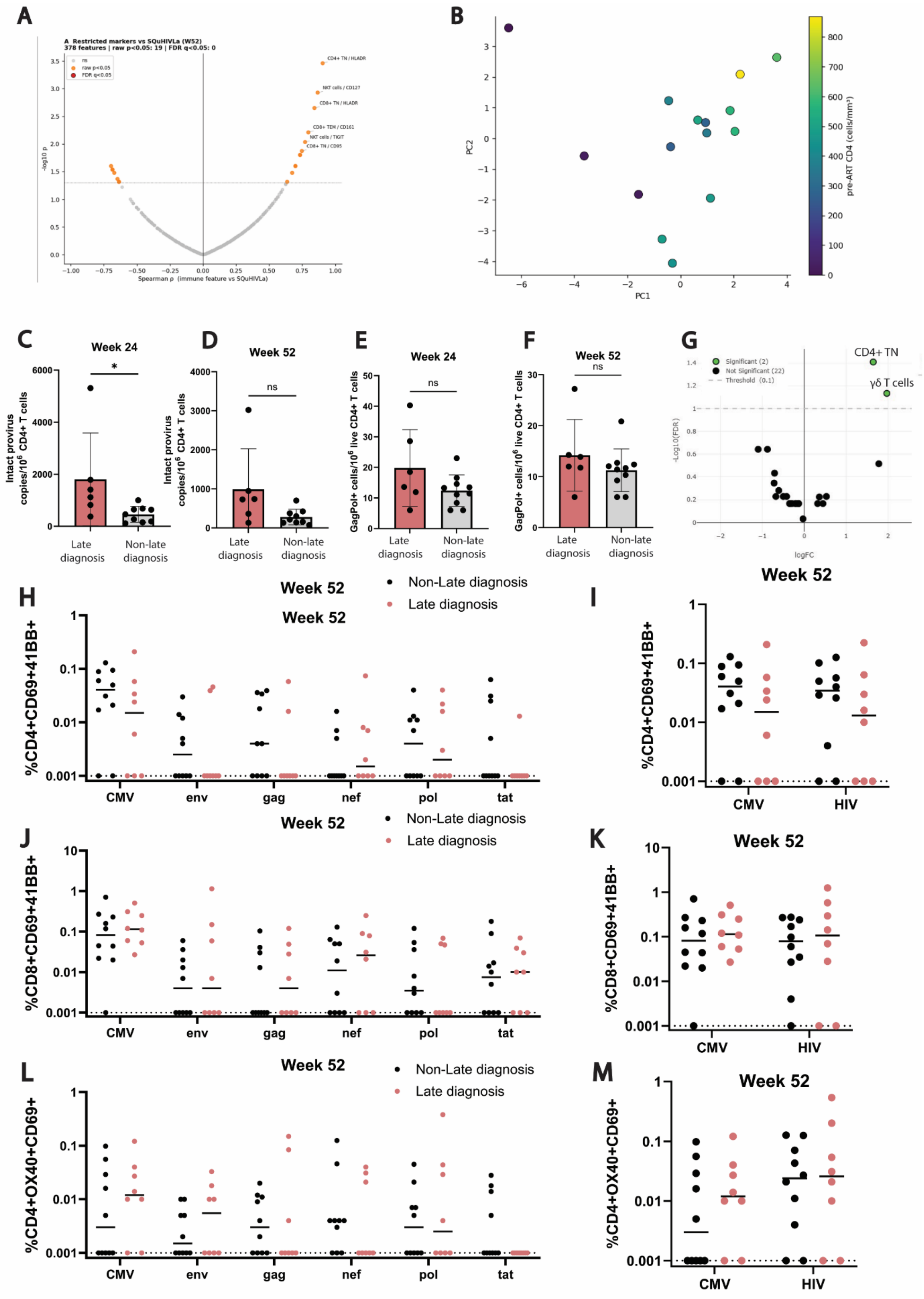

A
A Restricted markers vs SQuHIVLa (W52)
378 features | raw p<0.05: 19 | FDR q<0.05: 0
ns
raw p<0.05
FDR q<0.05
CD4+ TN / HLADR
NKT cells / CD127
CD8+ TN / HLADR
CD8+ TEM / CD161
NKT cells / TIGIT
CD8+ TN / CD95
-log10 p
Spearman ρ (immune feature vs SQuHIVLa)
B
PC1
PC2
pre-ART CD4 (cells/mm³)
C
Week 24
Intact provirus copies/10^6 CD4+ T cells
Late diagnosis
Non-late diagnosis
D
Week 52
ns
E
Week 24
GagPol+ cells/10^6 live CD4+ T cells
F
Week 52
G
CD4+ TN
γδ T cells
Significant (2)
Not Significant (22)
Threshold (0.1)
-Log10(FDR)
logFC
H
Week 52
Non-Late diagnosis
Late diagnosis
%CD4+CD69+41BB+
CMV
env
gag
nef
pol
tat
I
Week 52
HIV
J
Week 52
%CD8+CD69+41BB+
K
Week 52
L
Week 52
%CD4+OX40+CD69+
M
Week 52

**Fig S7. A)** Correlation of SQuHIVLa (MS expressing cells/$10^6$ CD4+ T cells) with all immune features (MFI and abundance) tested at the same time with one BH-FDR correction across all features (n = 10, FDR 0.05). **B)** PCA with all immune features, on Z-axis pre-ART CD4+ T cell count (cells/$mm^3$). **C-D)** Intact proviral reservoir at week 24 in late (n=6) and non-late (n=9) diagnosis group shown as Intact provirus copies/$10^6$ CD4+ T cells measured by IPDA at **C)** week 24 (p = 0.012) **D)** week 52 (p = 0.10) **E-F)** Inducible reservoir at week 24 in late (n = 6) and non-late (n = 10) diagnosis group shown as GagPol RNA expressing cells/$10^6$ CD4+ T cells measured by FISH-Flow after 16 hours of PMA/ionomycin stimulation at **E)** Week 24 (p = 0.12) **F)** Week 52 (p = 0.26) **G)** Volcano plot of significant differences in PBMC cluster abundance between late (n=5) and non-late (n=10) diagnosis groups at week 52 after ART initiation. Green dots are significant (FDR: 0.1). **H-M)** T cells after stimulation with HIV antigen (AIM), re stratified in late (pink) and non-late (black) diagnosis group (not significant, multiple unpaired T-tests with Welch correction for unequal sample sizes) percentages are normalized to DMSO-treated control. Dashed line indicates threshold of detection **H)** % CD4+ CD69+ CD4-1BB T cells after individual peptide stimulations **I)** sum of CD4+CD69+4-1BB+ T cell percentages of all HIV-specific peptide stimulations. **J) %** CD8+ CD69+ CD4-1BB+ T cells after individual peptide stimulations **K)** sum of CD8+ CD69+ CD4-1BB+ T cell percentages of all HIV-specific peptide stimulations **L)** % CD4+ OX40+ CD69+ T cells after individual peptide stimulations **M)** sum of CD4+ OX40+ CD69+ T cell percentages of all HIV-specific peptide stimulations.

**Novel IPDA primers and probes**

Given that the original IPDA primers published by Bruner *et al.* exhibited a failure rate of approximately 30% of samples, we redesigned the primers and probes to match 85% of full-length HIV-1 subtype B genomes available on the Lost Alamos National Library HIV Sequence Database (hiv.lanl.gov)[35, 36]. Newly designed primers and probes for IPDA and RRP30 quantification can be found in Table S4.

**IPDA validation**

Our novel primers and probes were first tested for specificity. The false-positive detection rate, tested using DNA from three negative donor PBMC samples analyzed in triplicates, was 0% for intact HIV-1 proviral DNA copies and 0.01% for the PSI or ENV regions. The linearity and sensitivity of the adapted IPDA were further validated using the pNL4.3 plasmid, demonstrating a lower limit of quantification of one copy (Fig. S1A). At input levels above 10,000 copies, oversaturation of the microchip was observed. Finally, the robustness of our assay was confirmed by successful quantification of intact, defective, and total HIV reservoirs in the participants analyzed.

**Reservoir quantification**

Not all reservoir quantification assays could be performed for all participants because of limited sample availability and subtype constraints. SQuHIVLa is available for subtype B and subtype C, the IPDA primers are subtype B specific, whereas FISH-Flow is less subtype-specific since detection is based on 40 probes and mismatches do not directly inhibit binding. In addition to subtype-specificity, sample availability was limited for a subset of participants. All reservoir quantification techniques were performed on all possible participants with the right subtype and enough sample availability (Table S2, S3).

**Clustering of cell populations**

Due to a machine error during running of the samples, the week 0 timepoint for one participant had to be excluded from further immunophenotyping analysis. Data cleanup was performed using the flowAI algorithm integrated in the OMIQ software. Downstream processing included scaling adjustments, equal sub-sampling, and manual gating to identify live single-cell populations (figure S2C-D). Subsampling and further analysis was performed both on full PBMCs as well as specifically on CD3+ T cells (Fig. S2D). High-dimensional analysis was conducted using the FlowSOM algorithm for clustering and UMAP for dimensionality reduction and two-dimensional visualization (26, 27). The number of clusters for FlowSOM clustering was set at 20 based on elbow metaclustering (Fig, S2E). After FlowSOM clustering, clusters 3, 4, 6, 7, 11 and 17 were manually subdivided based on their separation in the UMAP visualization (Fig. S2F-K). CD8+ central memory T cells (TCM) were not clearly distinguished by the clustering software, likely due to the large variability between week 0 and week 52 impeding the formation of two separate clusters. This cluster was therefore manually separated (Fig S2J). The software determined another exhausted CD8+ TEM cluster (18), however this cluster was excluded from the analysis as it was heavily biased by one donor (chrono 27). For CD3 subsampling, the number of clusters was set at 19 based on the elbow metaclustering function (Fig. S3A) and visual inspection of the least overlap in the UMAP. Although cells were subsampled on CD3+ expression, 5 non-T cell clusters with relatively high CD3 expression for non-T cells were included. We excluded these clusters based on low expression of CD3 (when compared to T cells) (Figure S3B). In addition, clusters 4, 8 and 9 were manually separated based on UMAP-visualization (Figure S3C-E). Differences in cluster abundance between groups were assessed using the edgeR algorithm (FDR < 0.1), with results visualised in volcano plots. Differences in cluster marker expression were examined using Significance Analysis of Microarrays (SAM) analysis.

| **Characteristic** | **Non-late** N = 17[1] | **Late** N = 18[1] | **p-value**[2] | **Overall** N = 35[1] |
|---|---|---|---|---|
| AIDS Defining Illnesses | | | | |
| No ADI | 17 (100%) | 11 (61%) | 0.008 | 28 (80%) |
| Wasting Syndrome | 0 (0%) | 5 (28%) | 0.045 | 5 (14%) |
| Pneumocystis Jirovecii Pneumonia | 0 (0%) | 4 (22%) | 0.10 | 4 (11%) |
| Mycobacterium Avium Complex | 0 (0%) | 1 (5.6%) | >0.9 | 1 (2.9%) |
| HIV Indicator Conditions | | | | |
| No Indicator Conditions | 5 (29%) | 2 (11%) | 0.2 | 7 (20%) |
| Sexually Transmitted Inections | 9 (53%) | 9 (50%) | 0.9 | 18 (51%) |
| Dysplasia | 0 (0%) | 1 (5.6%) | >0.9 | 1 (2.9%) |
| Herpes Zoster | 0 (0%) | 2 (11%) | 0.5 | 2 (5.7%) |
| Hepatitis B or C | 1 (5.9%) | 0 (0%) | 0.5 | 1 (2.9%) |
| Unexplained Leukocytopenia or thrombocytopania | 0 (0%) | 1 (5.6%) | >0.9 | 1 (2.9%) |
| Unexplained Fever | 0 (0%) | 1 (5.6%) | >0.9 | 1 (2.9%) |
| Severe or Atypical Psoriasis | 0 (0%) | 1 (5.6%) | >0.9 | 1 (2.9%) |
| Unexplained Weightloss | 2 (12%) | 2 (11%) | >0.9 | 4 (11%) |
| Unexplained Lymphadenopathy | 3 (18%) | 2 (11%) | 0.7 | 5 (14%) |
| Unexplained chronic diarrhoea | 1 (5.9%) | 2 (11%) | >0.9 | 3 (8.6%) |
| Hepatitis A | 0 (0%) | 1 (5.6%) | >0.9 | 1 (2.9%) |
| Community acquired pneumonia | 1 (5.9%) | 2 (11%) | >0.9 | 3 (8.6%) |
| Candidiasis | 0 (0%) | 3 (17%) | 0.2 | 3 (8.6%) |
| Co-Medication | | | | |
| Psychoanaleptics | 1 (7.1%) | 2 (12%) | >0.9 | 3 (9.7%) |
| Antivirals for systemic use | 1 (7.1%) | 2 (12%) | >0.9 | 3 (9.7%) |
| Antibacterials for systemic use | 8 (57%) | 12 (71%) | 0.5 | 20 (65%) |

| **Characteristic** | **Non-late** N = 17[1] | **Late** N = 18[1] | **p-value**[2] | **Overall** N = 35[1] |
|---|---|---|---|---|
| Psycholeptics | 4 (29%) | 8 (47%) | 0.3 | 12 (39%) |
| Immunosuppressants | 0 (0%) | 1 (5.9%) | >0.9 | 1 (3.2%) |
| Antimycotics for systemic use | 0 (0%) | 2 (12%) | 0.5 | 2 (6.5%) |
| Antiprotozoals | 0 (0%) | 2 (12%) | 0.5 | 2 (6.5%) |
| Corticosteroids for systemic use | 0 (0%) | 6 (35%) | 0.021 | 6 (19%) |
| Antimycobacterials | 0 (0%) | 2 (12%) | 0.5 | 2 (6.5%) |
| Sex hormones | 0 (0%) | 3 (18%) | 0.2 | 3 (9.7%) |
| Antihistamines | 0 (0%) | 2 (12%) | 0.5 | 2 (6.5%) |
| Antineoplastic agents | 0 (0%) | 2 (12%) | 0.5 | 2 (6.5%) |
| Antiepileptics | 1 (7.1%) | 0 (0%) | 0.5 | 1 (3.2%) |
| No co-medication | 3 (21%) | 4 (24%) | >0.9 | 7 (23%) |

[1]n (%)

[2]Fisher's exact test

**Table S1:** Participant HIV indicator conditions and co-medication.

| Participant Id | Sex at birth | Age at diagnosis | AIDS indicator condition at diagnosis | CD4+ T-cells pre-ART (cells/mm3) | HIV RNA pre-ART (copies/mL) | Diagnosis | HIV-1 Subtype | SQUIVLA | AIM | FISHFLOW | Immunophetyping | IPDA |
|---|---|---|---|---|---|---|---|---|---|---|---|---|
| CHRONO-01 | Female | 54 | Yes | 20 | 1540000 | Late | CRF06_cpx | | | X | X | |
| CHRONO-02 | Female | 32 | No | 870 | 6830 | Non late | B | X | X | X | X | X |
| CHRONO-03 | Male | 51 | Yes | 90 | 3260000 | Late | B | X | X | | | X |
| CHRONO-04 | Male | 47 | No | 90 | 22500 | Late | B | | | | | |
| CHRONO-05 | Male | 29 | No | 310 | 50000 | Late | B | | | | | |
| CHRONO-06 | Male | 41 | No | 570 | 19000 | Non late | F | | | X | X | |
| CHRONO-07 | Male | 46 | No | 400 | 157000 | Non late | B | X | X | | | X |
| CHRONO-08 | Male | 27 | No | 360 | 2200000 | Non late | B | X | X | X | X | X |
| CHRONO-09 | Male | 56 | No | 240 | 265000 | Late | B | | X | | | |
| CHRONO-10 | Male | 49 | No | 440 | 2370000 | Non late | B | X | X | X | X | X |
| CHRONO-11 | Male | 29 | No | 270 | 225000 | Late | B | X | X | X | X | X |
| CHRONO-12 | Male | 25 | No | 880 | 6990000 | Non late | CRF02_AG | | | | | |
| CHRONO-13 | Male | 56 | No | 470 | 215000 | Non late | B | X | X | X | X | X |
| CHRONO-14 | Male | 23 | No | 210 | 45800 | Late | B | | | | | |
| CHRONO-15 | Female | 41 | Yes | 30 | 71500 | Late | C | X | | X | X | |
| CHRONO-16 | Male | 61 | No | 480 | 114000 | Non late | B | X | X | X | X | X |
| CHRONO-17 | Male | 30 | Yes | 480 | 20800 | Late | B | X | X | | | X |
| CHRONO-18 | Male | 50 | No | 220 | 688000 | Late | CRF01_AE | | | | | |
| CHRONO-19 | Male | 39 | No | 570 | 4700 | Non late | F | | | X | X | |
| CHRONO-20 | Male | 69 | No | 1030 | 44800 | Non late | B | X | X | | | X |
| CHRONO-21 | Male | 36 | No | 380 | 4010 | Non late | B | | | | | |
| CHRONO-22 | Male | 24 | No | 630 | 6010 | Non late | B | X | X | X | X | X |
| CHRONO-23 | Male | 24 | No | 420 | 1130000 | Non late | CRF02_AG | | | | | |
| CHRONO-24 | Male | 51 | No | 250 | 445000 | Late | B | X | X | | | X |
| CHRONO-25 | Male | 26 | No | 1070 | 423 | Non late | B | | X | | | |
| CHRONO-26 | Male | 38 | No | 630 | 8750 | Non late | CRF01_AE | | | | | |
| CHRONO-27 | Female | 39 | Yes | 0 | 2210000 | Late | CRF02_AG | | | X | X | |
| CHRONO-28 | Male | 60 | No | 210 | 26200 | Late | B | X | X | | | |
| CHRONO-29 | Male | 67 | No | 210 | 315000 | Late | B | | | | | |
| CHRONO-30 | Female | 48 | Yes | 20 | 750000 | Late | B | X | X | X | | X |
| CHRONO-31 | Male | 66 | Yes | 230 | 3350000 | Late | B | Excl* | Excl* | Excl* | Excl* | Excl* |
| CHRONO-32 | Male | 31 | No | 510 | 106000 | Non late | B | X | X | X | X | X |
| CHRONO-33 | Male | 36 | No | 260 | 81900 | Late | B | X | X | X | X | X |
| CHRONO-34 | Male | 31 | No | 390 | 73800 | Non late | CRF02_AG | | | X | X | |
| CHRONO-35 | Male | 26 | No | 330 | 122000 | Late | C | | | | | |

**Table S2:** individual participant clinical characteristics and inclusion in assays. *excluded because the viral load was not suppressed.

| Characteristic | Total N = 35[1] | IPDA N = 15[1] | FISH-Flow N = 16[1] | SQuHIVLa N = 17[1] | AIM N = 18[1] | Immune profiling N = 15[1] |
|---|---|---|---|---|---|---|
| Sex assigned at birth | | | | | | |
| Male | 30 (86%) | 13 (87%) | 11 (69%) | 14 (82%) | 16 (89%) | 11 (73%) |
| Female | 5 (14%) | 2 (13%) | 5 (31%) | 3 (18%) | 2 (11%) | 4 (27%) |
| Age at diagnosis | 39 (29, 51) | 46 (30, 51) | 39 (31, 49) | 46 (31, 51) | 47 (30, 56) | 39 (31, 49) |
| HIV-1 subtype | | | | | | |
| B | 24 (69%) | 15 (100%) | 10 (63%) | 16 (94%) | 18 (100%) | 9 (60%) |
| C | 2 (5.7%) | | 1 (6.3%) | 1 (5.9%) | | 1 (6.7%) |
| CRF01_AE | 2 (5.7%) | | | | | |
| CRF02_AG | 4 (11%) | | 2 (13%) | | | 2 (13%) |
| CRF06_cpx | 1 (2.9%) | | 1 (6.3%) | | | 1 (6.7%) |
| F | 2 (5.7%) | | 2 (13%) | | | 2 (13%) |
| Diagnosis | | | | | | |
| Late | 18 (51%) | 6 (40%) | 6 (38%) | 8 (47%) | 8 (44%) | 5 (33%) |
| Non-Late | 17 (49%) | 9 (60%) | 10 (63%) | 9 (53%) | 10 (56%) | 10 (67%) |
| CD4+ T cell pre-ART (cells/mm3) | 360 (210, 510) | 440 (260, 510) | 415 (145, 540) | 400 (250, 480) | 420 (250, 510) | 440 (260, 570) |
| Viral load pre-ART (copies/mL) | 114,000 (22,500, 750,000) | 157,000 (44,800, 750,000) | 110,000 (45,250, 1,145,000) | 114,000 (44,800, 445,000) | 135,500 (26,200, 445,000) | 106,000 (19,000, 1,540,000) |
| CD4+ T cell week 0 (cells/mm3) | 390 (270, 680) | 500 (270, 680) | 545 (170, 680) | 500 (270, 680) | 510 (340, 680) | 570 (270, 680) |
| Viral load week 0 (copies/mL) | 14,000 (795, 46,500) | 14,600 (1,250, 74,600) | 10,090 (1,565, 80,250) | 4,780 (1,250, 44,900) | 9,690 (795, 44,900) | 15,400 (1,250, 85,900) |
| Median time to viral suppression* (days) | 37 (28, 90) | 86 (31, 125) | 35 (28, 89) | 48 (34, 91) | 45 (31, 91) | 34 (27, 87) |
| NA | 1 | | | | | |

[1]n (%); Median (Q1, Q3). ART: antiretroviral therapy. *<30 copies/mL.

**Table S3:** Participant characteristics in each assay.

| | |
|---|---|
| **Primers and Probes IPDA HIV-1 Subtype B** | |
| IPDA_B_PSI_FWD | 5' GAA-CAG-GGA-CYY-GAA-AGC-GAA-AG 3' |
| IPDA_B_PSI_RVS | 5' CCC-ATC-KCT-CTC-CTT-CTA-GCC-TC 3' |
| IPDA_B_ENV_RVS_1 | 5' CGC-CTC-AAT-AGC-CCT-CAG-C 3' |
| IPDA_B_ENV_RVS_2 | 5' GCG-CCT-CAA-TAG-CCT-TCA-GC 3' |
| IPDA_B_ENV_RVS_3 | 5' GCG-CCT-CAA-TAG-CTC-TCA-GC 3' |
| IPDA_B_ENV_FWD_1 | 5' GAG-GGA-CAA-TTG-GAG-AAG-TGA-ATT-ATA-TAA-ATA 3' |
| IPDA_B_ENV_FWD_2 | 5' TGA-AAG-ACA-ATT-GGA-GAA-GTG-AAC-TAT-ATA-AAT-A 3' |
| IPDA_B_ENV_FWD_3 | 5' ATG-AGG-GAT-AAT-TGG-AGA-AGT-GAA-TTA-TAT-AAA-TA 3' |
| IPDA_B_ENV_Probe_HEX | 5' HEX - CAG-CAG-GAA-GCA-CTA-TGG-GCG-CA - BHQ®-1 3' |
| IPDA_B_PSI_Probe_FAM | 5' 6-FAM - ACG-CAG-GAC-TCG-GCT-TGC-TGA-AG - BHQ®-1 3' |
| IPDA_B_ENV_A3GProbe | 5' CAG-CAA-AAA-GCA-CTA-TAG-GCG-CA 3' |
| **Primers and Probes RPP30 quantification** | |
| RPP30_FWD | 5' AGA-TTT-GGA-CCT-GCG-AGC-G 3' |
| RPP30_RVS | 5' GTG-AGC-GGC-TGT-CTC-CAC 3' |
| RPP30_Shear_FWD | 5' CCA-TTT-GCT-GCT-CCT-TGG-GA 3' |
| RPP330_Shear_RVS | 5' AGC-TGA-GAC-AAT-CAT-GCT-GGC 3' |
| RPP30_Probe | 5' HEX - CTG-ACC-TGA-AGG-CTC-TGC-GCG - BHQ®-1 3' |
| RPP30_Shear_Probe | 5' 6-FAM - CGG-CTT-CCT-CCT-TTG-CAT-GCT-CTG - BHQ®-1 3' |

**Table S4:** Adjusted primer sequences for IPDA

| **Characteristic** | **Late** N = 18[1] | **Non-Late** N = 17[1] | **p-value**[2] | **Overall** N = 35[1] |
|---|---|---|---|---|
| **Sex assigned at birth** | | | 0.3 | |
| Male | 14 (78%) | 16 (94%) | | 30 (86%) |
| Female | 4 (22%) | 1 (5.9%) | | 5 (14%) |
| **Age at diagnosis** | 48 (30, 54) | 36 (27, 46) | 0.2 | 39 (29, 51) |
| **Mode of transmission** | | | 0.6 | |
| MSM | 9 (50%) | 12 (71%) | | 21 (60%) |
| Heterosexual | 5 (28%) | 3 (18%) | | 8 (23%) |
| IV drugs and MSM | 1 (5.6%) | 0 (0%) | | 1 (3%) |
| Unknown | 3 (17%) | 2 (12%) | | 5 (14%) |
| **HIV-1 subtype** | | | 0.6 | |
| B | 13 (72%) | 11 (65%) | | 24 (69%) |
| Non-B | 5 (28%) | 6 (35%) | | 11 (31%) |
| **Region of birth** | | | 0.7 | |
| Africa | 0 (0%) | 2 (12%) | | 2 (6%) |
| Americas | 2 (11%) | 1 (5.9%) | | 3 (9%) |
| Asia | 2 (11%) | 1 (5.9%) | | 3 (9%) |
| Europe | 14 (78%) | 13 (76%) | | 27 (77%) |
| **Initial ART regimen** | | | 0.14 | |
| TDF/FTC+DTG | 14 (78%) | 9 (53%) | | 23 (66%) |
| ABC/3TC/DTG | 3 (17%) | 6 (35%) | | 9 (26%) |
| TAF/FTC+DTG | 1 (5.6%) | 0 (0%) | | 1 (3%) |
| TAF/FTC/BIC | 0 (0%) | 2 (12%) | | 2 (6%) |
| **ART regimen at week 52** | | | 0.11 | |
| TDF/FTC+DTG | 5 (28%) | 2 (12%) | | 7 (20%) |
| ABC/3TC/DTG | 0 (0%) | 4 (24%) | | 4 (11%) |
| TAF/FTC+DTG | 3 (17%) | 0 (0%) | | 3 (9%) |
| TAF/FTC/BIC | 1 (5.6%) | 3 (18%) | | 4 (11%) |
| DTG/3TC | 3 (17%) | 5 (29%) | | 8 (23%) |
| TDF/3TC/DOR | 1 (5.6%) | 1 (5.9%) | | 2 (6%) |
| CAB+RPV | 1 (5.6%) | 0 (0%) | | 1 (3%) |
| Drop-out | 4 (22%) | 2 (12%) | | 6 (17%) |
| **CD4+ T-cell count pre-ART (cells/mm3)** | 215 (90, 260) | 510 (420, 630) | <0.001 | 360 (210, 510) |
| **CD4+ T-cell count week 24 (cells/mm3)** | 350 (200, 500) | 890 (780, 1,160) | <0.001 | 530 (340, 880) |
| Missing | 1 | 3 | | 4 |
| **CD4+ T-cell count week 52 (cells/mm3)** | 370 (200, 540) | 810 (640, 1,120) | <0.001 | 620 (410, 810) |

| **Characteristic** | **Late** N = 18[1] | **Non-Late** N = 17[1] | **p-value**[2] | **Overall** N = 35[1] |
|---|---|---|---|---|
| Missing | 4 | 2 | | 6 |
| **Viral load pre-ART (copies/mL)** | 245,000 (50,000, 750,000) | 73,800 (6,830, 215,000) | 0.10 | 114,000 (22,500, 750,000) |
| **Median time to viral suppression <30 copies/mL (days)** | 48 (35, 90) | 31 (28, 89) | 0.3 | 37 (28, 90) |
| Not suppressed | 1 | 0 | | 1 |

[1]n (%); Median (Q1, Q3)

[2]Fisher's exact test; Wilcoxon rank sum exact test; Pearson's Chi-squared test

**Table S5:** Overall participant characteristics in the cohort. Data are n (%) or median (IQR). MSM: men who have sex with men; ART: antiretroviral therapy; TDF: tenofovir disoproxil fumarate; FTC: emtricitabine; DTG: dolutegravir; ABC: abacavir; 3TC: lamivudine; TAF: tenofovir alefenamide; BIC: bictegravir; DOR: doravirine; CAB: cabotegravir; RPV: rilpivirine.

| Cell type | Markers |
| --- | --- |
| **CD8+ TCM** | CD3+CD8+CD45RA-CCR7+CD28+CD27+ |
| **CD8+ TEM** | CD3+CD8+CD45RA-CCR7- |
| **CD8+ TEMRA** | CD3+CD8+CD45RA+CCR7-CD28-CD27- |
| **CD8+ TN** | CD3+CD8+CD45RA+CCR7+CD28+CD27+ CD95- |
| **CD4+ TCM** | CD3+CD4+CD45RA-CCR7+CD28+CD27+ |
| **CD4+ TN** | CD3+CD4+CD45RA+CCR7+CD28+CD27+ CD95- |
| **CD4+ TEM (CD57+ CD25-)** | CD3+CD4+CD45RA-CCR7-CD28-CD27- CD57+ CD25- |
| **CD4+ Treg** | CD3+CD4+CD127dim CD25++ CD27+ CD28+ CD95+ |
| **gdTCR** | CD3+gdTCR+CD4-CD8- |
| **Naïve B-cells** | CD19+ CD20+IgD+IGM+ CD27- |
| **Mature (IgG+) B-cells** | CD19+CD20+HLA-DR+IgM-IgD-IgG+CD27+ |
| **IgD+ IgM- B-cells** | CD19+CD20+IgD+IgM-IgG-CD27- |
| **DCs** | CD3-CD19-CD56-CD14-HLADR+CD123-CD11c+ |
| **pDCs** | CD3-CD19-CD56-CD14-HLADR+CD123+ |
| **Conventional DCs** | CD3-CD19-CD56-CD14-HLADR+CD123-CD11c+ |
| **DR- Basophils** | CD123+CD19-CD20-HLADR-CD56-CD16-CD11c- |
| **Mature NKs** | CD3-CD14-CD20-CD56+CD16+ |
| **Classical monocytes** | HLADR+CD11b+CD14+CD16- |
| **Intermediate monocytes** | HLADR+CD11b+CD14+CD16+ |
| **Non-classical monocytes** | HLADR+CD11b+CD14+CD16++ |
| **MAIT** | CD4-CD8-CD3+CD161+KLRG1+CCR5+CD127+ |

**Table S6:** annotation of clusters based on marker expression
